# Fixing reference errors efficiently improves sequencing results

**DOI:** 10.1101/2022.07.18.500506

**Authors:** S. Behera, J. LeFaive, P. Orchard, M. Mahmoud, L. F. Paulin, J. Farek, D. C. Soto, S.C.J. Parker, A. V. Smith, M. Y. Dennis, J. M. Zook, F.J. Sedlazeck

## Abstract

The GRCh38 reference is the current standard in human genomics research and clinical applications, but includes errors across 33 protein-coding genes, including 12 with medical relevance. Current studies rely on the correctness of this reference genome and require an accurate and cost-effective way to improve variant calling and expression analysis across these erroneous loci. We identified likely artifacts in GTEx, gnomAD, 1000 Genomes Project, and other important genomic resources leading to wrong interpretations for these genes. Here, we present FixItFelix together with a modified GRCh38 version that improves the subsequent analysis across these genes within minutes for an existing BAM/CRAM file. We showcase these improvements over multi-ethnic control samples across short and long-read DNA-, and RNA-sequencing. Furthermore, applying our approach across thousands of genomes demonstrates improvements for population variant calling as well as eQTL studies. Still, some genes e.g., *DUSP22* indicate mixed results due to their complexity.

## Introduction

The identification of genetic variation in individuals and populations is essential for all genomic analyses to answer questions related to evolution, diversity, diseases, and biological processes in general^1,2,3^. To identify variation, sequences are typically mapped to a reference genome^4,5^, though *de novo* genome assembly approaches using long reads are advancing rapidly^6,7^. Typically, in both cases, methods compare to a single reference genome to form a unified coordinate system that enables the comparison of differences across multiple projects and thus enables novel insights. For humans, we have had a reference genome since the Human Genome Project released its first version in 2001^8^. Since then multiple updates have been made with the current version (GRCh38) slowly being adopted over the last decade^9^. Most recently, the Telomere-to-Telomere (T2T) Consortium released a new complete human genome reference (T2T-CHM13)^10^. Though initial studies show promising results^11^, T2T-CHM13 currently lacks many of the resources and annotations that exist for GRCh38 that likely hindering the community’s transition to this new reference.

Over the past years, it has become clear that a reference genome cannot be a single best representation of the human population or any given species, but it eases the comparison across studies and holds the promise of curation resources^12^. As such, the human reference genome never included all major alleles or all minor alleles from a population, but rather a mix of haplotypes from multiple individuals^9^. Nevertheless, multiple updates have corrected errors, added alternate loci, and improved certain regions of the genome to make the representation more complete and, thus, improve variant calling and comparison. Most recently, our work identified remaining issues with the most commonly used reference genomes (GRCh37+38), where certain regions of the genome have been duplicated along chromosome 22^13^. These include at least three medically relevant genes for inherited diseases, as well as one relevant for somatic variant calling^14^. Continuing this work together with the T2T group revealed other artifacts along GRCh38, including additional false duplications and missing copies (or collapses) of segmental duplications^11^. As these regions also impact medically relevant genes, we are eager to correct them and improve mapping and variant calling.

In this work, we focus on providing a solution for these reference issues to improve variant calling across 33 protein-coding genes including 12 medically relevant genes. To achieve this, we propose a modified GRCh38 reference that includes several masked regions as well as newly introduced decoy contigs. Using this reference, we demonstrate improvements in mapping and single-nucleotide variant (SNV) calling across different ancestries. In addition, we propose a rapid, localized remapping framework that improves the alignment of short- or long-reads across targeted regions and provides a modified alignment (BAM/CRAM) file that can be used for subsequent variant identification. This approach improves accuracy not only for human individuals of European ancestry, HG002 (Genome in a Bottle [GIAB] NIST Reference Material)^15^, but also for more ancestrally diverse groups. Furthermore, we assess the benefits not only for whole-genome and exome sequencing of short and long reads but also for RNA sequencing analysis, making this an important analytical change for many studies to come. Maybe even more importantly, we highlight its improvements across different human ancestries such as African, European and Asian populations. Thus, we show clear improvements for these genes across the 3,202 samples from the 1000 Genomes Project (1KGP)^16^ discovering novel alleles across these important genes and regions. We further investigate if these SNV improvements have a significant impact on phenotypic traits. Lastly, we highlight the importance of these introduced changes by showing improvements for eQTL studies. All together, we demonstrate the importance of the newly suggested changes to GRCh38 itself and a computationally efficient solution to improve existing mapped genomic (BAM/CRAM) data.

## Results

### Identification of GRCh38 errors

From our previous work^11^, we detected errors in the GRCh38 reference genome, including i) 1.2 Mbp of falsely duplicated regions: regions of the genome that were present more often than they should be, and ii) 8.04 Mbp of falsely collapsed regions: regions with paralogs missing in the reference. **Figure 1A** describes the issues with these regions that can lead to incorrect mapping of reads and subsequent biases in analysis. This has potential implications across current published work as these regions include medically relevant genes that have been reported through, e.g., GTEx and GWAS studies^17,18^. To improve mapping and variant calling, we generated a modified GRCh38 reference by first masking out the 1.2 Mbp of extra copies of falsely duplicated regions. From the collapsed regions, we selected a targeted set of three medically relevant genes (*MAP2K3, KCNJ18*, and *FANCD2*), two human-specific duplicated genes (*GPRIN2* and *DUSP22*), and their homologous genes and pseudogenes. For the GRCh38 missing genes, we used the T2T-CHM13 reference genome, which was found to correct all identified duplication errors^10^. Specifically, we identified all duplicate homologs in T2T-CHM13 syntenic to the “collapsed” region in GRCh38 and by comparing the references, narrowed in on genomic loci not represented in GRCh38. We then added these missing sequences as decoys to our modified GRCh38 reference genome. See **Methods** for details.

**Figure 1:**
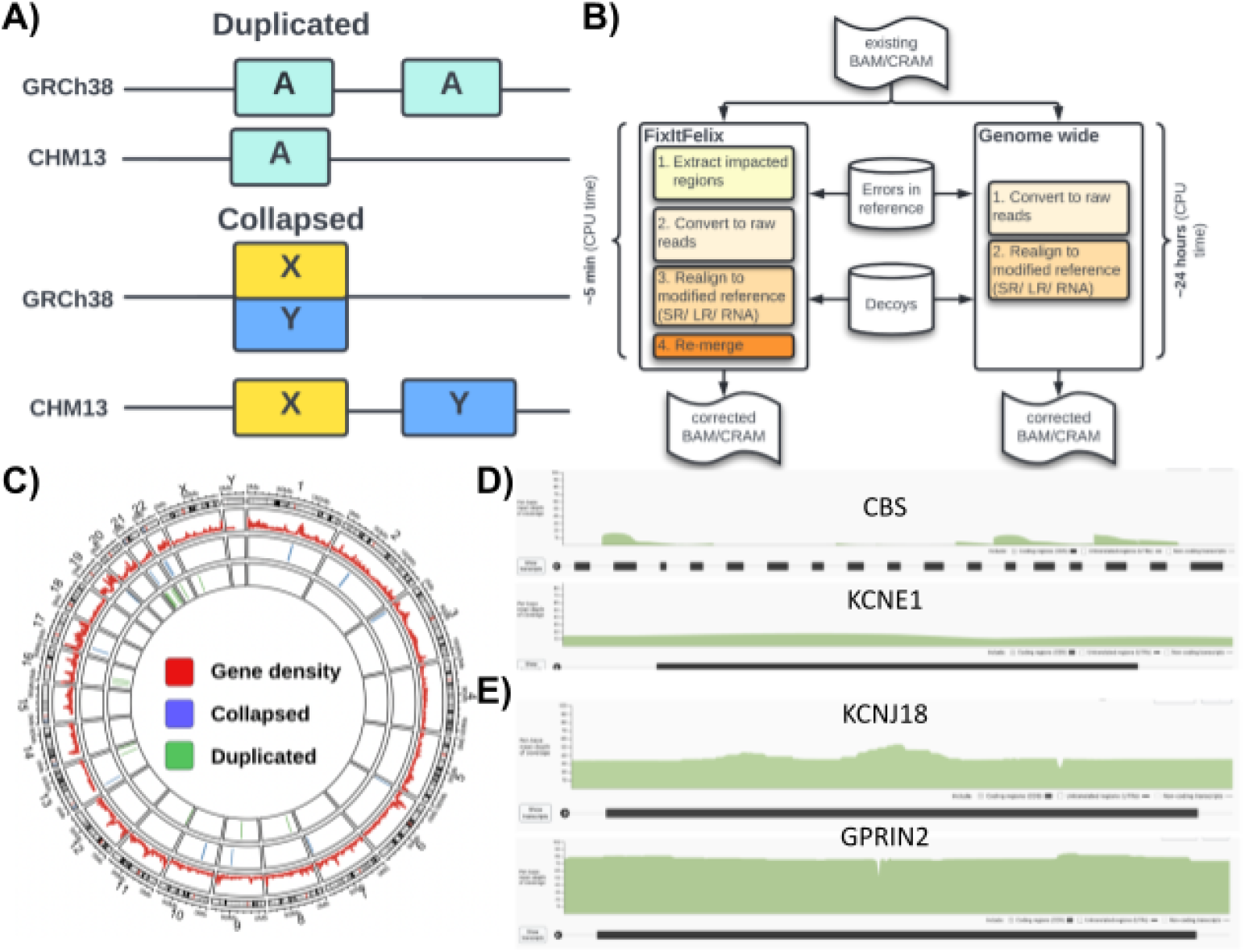
Falsely duplicated and collapsed regions of GRCh38 reference. **A)** Cartoon showing the duplicated and collapsed errors. For duplicated errors, one extra copy that is absent in T2T-CHM13 reference is present in GRCh38. For collapsed regions, the two separate copies are merged into one region. **B)** Our pipeline (FixItFelix), shown in the left side of the figure, extracts the sequences from impacted regions and remaps them to the modified reference for subsequent analysis. This takes only 5~8 minutes as compared to >24 hour remapping of all sequences to the modified reference genome i.e. global realignment (shown in the right side). **C)** 12 wrongly duplicated regions that include 22 protein-coding genes, 18 pseudogenes and total 2,032,012 bp (1,021,203 bp of correct regions, and 1,010,809 bp of false duplication); 9 collapsed regions that include 9 genes and **total** 843,139 bp. **D)** gnomAD track showing lower than normal whole genome sequencing (WGS) coverage of falsely duplicated genes (CBS & KCNE1) **E)** gnomAD track showing higher than normal WGS coverage of collapsed genes KCNJ18 and GPRIN2 where one or more paralogs are missing.

Using this modified reference with masking and additional decoys, we developed a new approach (FixItFelix) (https://github.com/srbehera/FixItFelix) to efficiently re-align only the affected reads to the modified GRCh38, correcting existing BAM/CRAM files and subsequent variant calling. FixItFelix is open source (MIT license) and has different modules for short-read, long-read DNA and RNA sequencing reads. Using FixItFelix, a ~30x genome coverage bam file can be corrected with the new reference in around 5~8 minutes CPU time, whereas traditional remapping often takes ~24 CPU hours. The left panel of **Figure 1B** shows an outline of FixItFelix. To ensure that we captured potential mis-mappings, we collected reads originally mapping to any region homologous to the falsely duplicated and falsely collapsed regions, and re-mapped them to our modified GRCh38 reference. We also tested FixItFelix with whole genome mapping (the right panel of **Figure 1B**), to ensure our novel approach appropriately called variants and we find results are concordant. **Figure 1C** shows the location and the number of regions that are impacted by issues along the GRCh38 reference genome. **Supplement Table 1 & 2** contains the coordinates of falsely duplicated regions and falsely collapsed regions and the genes that are present in these regions. These genes have problematic coverage in large studies like gnomAD that use GRCh38, with lower than expected coverage for falsely duplicated genes like *CBS* and *KCNE1* (as shown in **Figure 1D**), and higher than expected coverage in parts or all of collapsed genes like *KCNJ18* and *GPRIN2* (as shown in **Figure 1E**).

### Improving variant calling with modified GRCh38 based on GIAB

To measure if the modified GRCh38 reference improves mapping quality and variant calling, we first performed a series of experiments on the well-studied dataset of the HG002 sample. For benchmarking the variant calls using the modified reference genome, the reads were extracted from the original mapping (35x coverage and 2×150 bp Illumina short reads mapped to the original GRCh38 reference) and then remapped to the modified GRCh38 reference sequence.

As the falsely duplicated regions were masked in the modified reference sequence, it was expected that the mapping quality (MAPQ) would be improved. This is because the reads that were ambiguously aligned to duplicated regions would be mapped to a single region. In contrast, the mapping quality was expected to be reduced for collapsed regions as the reads could be mapped to two different regions i.e., the collapsed region and the decoy, instead of the one collapsed region. Our experiments showed that the mapping quality for falsely duplicated regions significantly improved with 78% fewer reads that were mapped to multiple locations (MAPQ = 0: 358,644 in original vs 103,392 in remapping) in the original mapping (Wilcoxon Rank Sum test *p*-value = 7.396e-07). **Figure 2A** shows the details per region. Conversely, the mapping quality analysis of collapsed regions showed that the number of reads mapping to different locations were increased by 20% (MAPQ = 0: 111,685 reads in original vs 192,280 reads in remapping) as compared to original mapping. Nevertheless, the average mapping quality was as expected, reduced as shown in **Figure 2B**. See **Supplement Table 3 & 4** for details.

**Figure 2:**
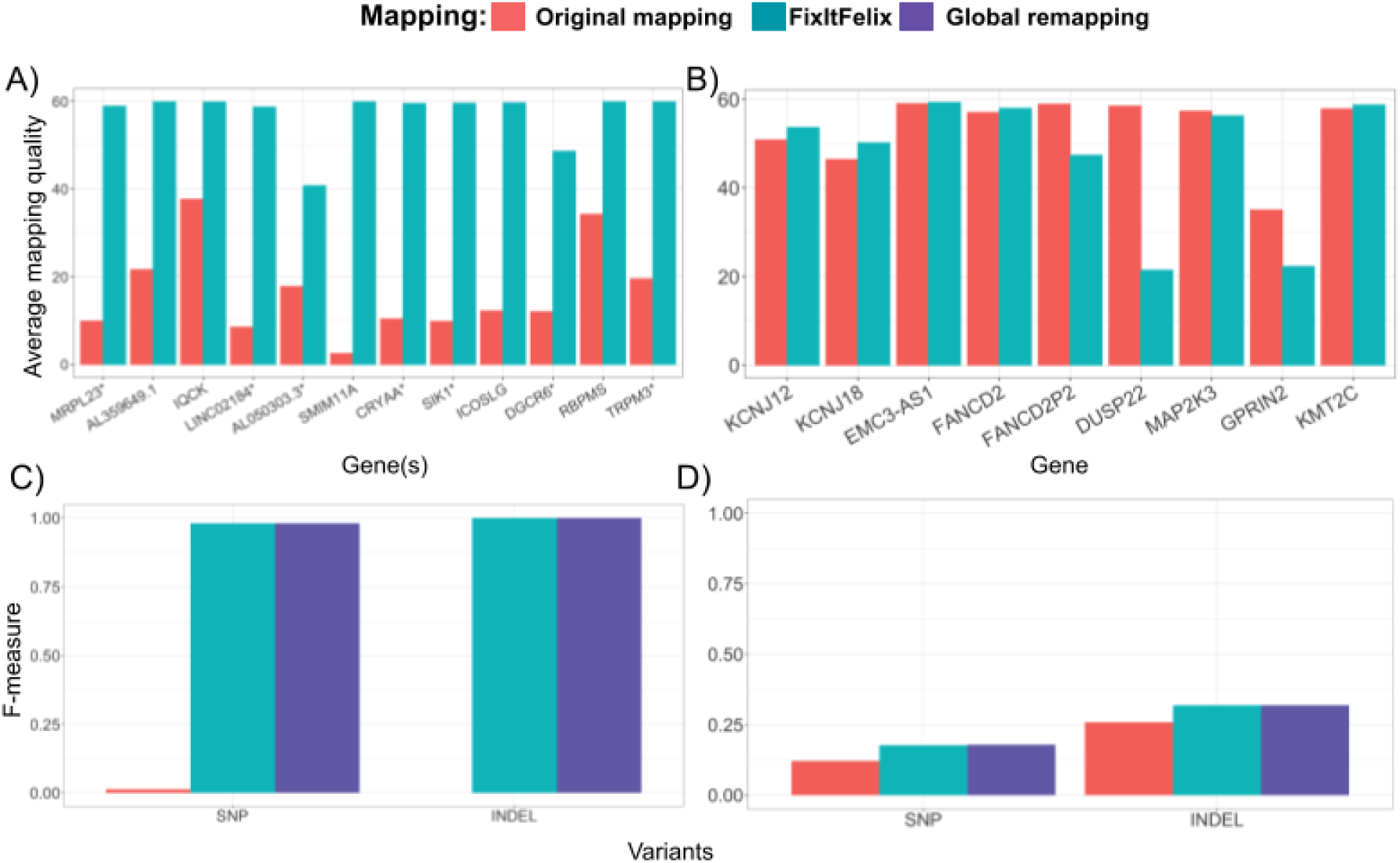
Performance improvements over modified GRCh38 via remapping. **A)** Improvements in mapping quality for duplicated regions **B)** Changes to mapping quality for collapsed regions. **C)** SNV and INDEL calling is improved over the mapping to the GRCh38 reference across six genes that are covered in CMRG benchmarking. Furthermore it clearly shows that a regional approach (FixItFelix) is concordant with global remapping. **D)** SNV & INDEL calling improvement for collapsed regions again highlighting that global alignment and regional alignment showing similar accuracy.

Given the changes in mapping, we next assessed the variant call performance using the benchmark set for challenging medically relevant genes (CMRG v1.0), which included five medically relevant genes (*KCNE1, CBS, CRYAA, TRAPPC10, DNMT3L)* affected by our masking of false duplications and one gene (*KMT2C)* affected by our decoys for collapsed duplications^13^. The GIAB developed this curated benchmark using *de novo* assembly, which identified and correctly resolved the falsely duplicated regions. For falsely duplicated regions, the short read BWA MEM-GATK^19^ variant calls for both SNV and INDEL greatly improved when the modified reference was used for re-mapping (**Figure 2C**). For SNVs, the recall score achieved by remapping was 1.0, which was improved significantly compared to the original mapping (0.007). Similarly, the precision and F-measures from the remapping (0.961 and 0.980 respectively) also improved significantly compared to the original mapping (0.063 and 0.012 respectively). For INDEL calls, the remapping was also able to call all 16 true variants, i.e. recall score 1.0 (original mapping: 0.0, no true INDEL calls) with the precision of 1.0 (original: 0.0). See **Supplement Table 5** for details. For the falsely collapsed regions, we observed that remapping produced mainly improved precision by removing false positive variants caused by mismapped reads (as shown in **Figure 2D**). For SNV calls for *KMT2C* only, the recall improved from 0.949 to 0.974 and the precision score from 0.064 to 0.098. For INDEL calls, remapping did not show any improvements for the *KMT2C gene* as compared to original mapping. However, there was an improvement in both the precision (0.152 to 0.195) and thus the F-measures (0.259 to 0.319). See **Supplement Table 6 for details.**

We have measured the outcomes by realigning the reads from the entire genome as well using FixItFelix. **Figure 2 C & D** show the concordance of the results.

### General improvements of variant calling

The CMRG benchmark dataset that was used for validation is limited by the number of genes it characterized. Thus, many improvements are not covered. Therefore, expanding on the approach used for CMRG, we utilized the phased HiFi assembly^6^ and dipcall^20^ (see **Methods**) and treated the resulting VCF and BED as a draft benchmark. Furthermore, we confirmed that GIAB CMRG and dipcall results were concordant across all regions included in CMRG (see **Supplement Table 7**).

The SNV and INDEL calls were clearly improved by the modified reference in falsely duplicated regions when benchmarking against the draft benchmark i.e., dipcall VCF and BED (as shown in **Figure 3A**). For SNVs, the improvement of variant calls is significant (Friedman rank sum test *p*-value = 0.02307) as the recall went up from 0.13 to 0.85 with the improved precision score. We also observed similar performance for INDEL calls. For collapsed regions, the precision and thus overall F-scores were improved from 0.22 and 0.32 to 0.452 and 0.453 respectively. Thus, we showed that across more genes, not covered in CMRG dataset, the proposed reference modifications are showing an overall improvement for mapping and variant calling.

**Figure 3:**
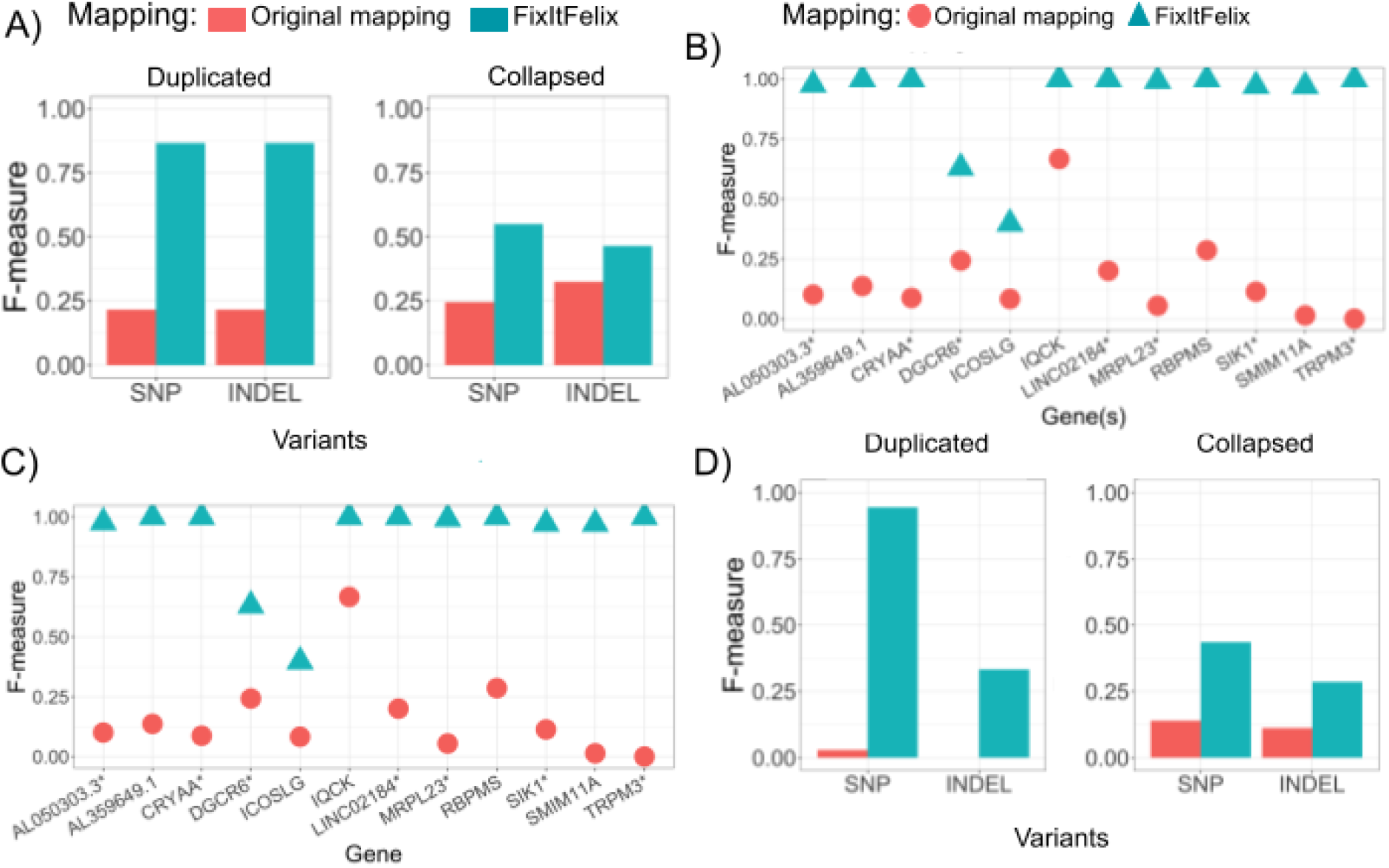
SNV benchmarking on HG002 using dipcall benchmark set: **A)** Benchmarking of SNV and INDEL using GATK variant calls with original mapping (original GRCh38 reference genome) and FixItFelix (modified GRCh38 reference). The region-wise (labeled by gene(s) inside the region) benchmarking are shown for **B)** duplicated regions, where more than one gene were impacted at some regions and **C)** collapsed regions, where each region contains only one gene. **D)** Benchmarking of whole-exome sequencing datasets that shows the improvement of variant calling on exon regions when the modified reference was used.

To determine if variant accuracy differs between affected regions, we analyzed the variant call performance for each individual region impacted by falsely duplicated and collapsed events. For falsely duplicated regions, we observed a clear improvement of SNV calls by using modified references over the original. The F-measures were 1.0 for 6 out of 12 regions and 0.97~0.99 for other 4 regions (see **Figure 3B)**. The recall, precision and F-measure scores for all regions are shown in **Supplement Tables 8 & 9.** The two regions with lower performance had additional true segmental duplications that caused mapping challenges even after masking the falsely duplicated sequence. Collapsed duplication showed more variable performance between regions. The modified reference substantially improved accuracy in the five regions containing genes *FANCD2, EMC3-AS1, KCNJ12, KCNJ18*, and *MAP2K3.* The *KMT2C* gene showed moderate improvements, but some sequences appeared to still be missing from GRCh38. However, two genes (*DUSP22* and *GPRIN2)* showed slightly lower F-measures and for pseudogene *FANCDP2*, it went down from 0.941 to 0.318. Upon curating alignments, the larger regions containing *FANCD2*, *DUSP22*, and *GPRIN2* appeared to have challenges both in making reliable assembly-based benchmarks and in mapping reads to GRCh38 with the decoys. GRCh38 may have structural errors in these genes in addition to the collapsed duplication, and common structural variation in the population may impact variant call accuracy. While most parts of the *FANCD2*, *DUSP22*, and *GPRIN2* genes affected by these decoys appear to be improved, these regions are more complicated and some regions in and around these genes may have performance reduced by the decoys.

We also compared the variant calling performance for both falsely duplicated and collapsed regions with the original and modified GRCh38 reference genome using long-read sequencing (PacBio HiFi and ONT) of HG002 sample (see **Methods**). Using the PacBio HiFi reads, we observed a slightly lower precision but with an overall improved F-score for SNV (from 0.251 to 0.308) and INDEL (0.421 to 0.452) calling as well for duplicated regions. We also obtained an improved precision and recall and thus overall F-score for SNV (0.242 to 0.327) and INDEL (0.451 to 0.564) calling across the collapsed regions. Similarly, for ONT reads we also observed a general improvement in SNV and INDEL calls. **Supplement Table 10** contains all the results for long read datasets.

Given the clear improvement across whole genome sequencing (long- and short-reads) data in most regions, we next investigated the effects of the modified reference to Whole Exome Sequencing (WES) and RNA sequencing. For WES-based SNV calls in duplicated regions, the use of the modified reference greatly improves the performance with a high recall score of 0.909 with the precision 0.984 and F-measure 0.945 as compared to the original reference based recall, precision and F-measures 0.015, 0.333 and 0.029, respectively (as shown in **Figure 3D**). For SNV calls in collapsed regions, we observed a slightly increased recall (0.389 vs 0.333) with the modified reference and a higher precision (0.632 vs 0.067). So, the F-measure improved from 0.159 to 0.472 (as shown in **Figure 3D**). There were only a few INDEL in the erroneous regions (nine in duplicated and three in collapsed regions) found in the dipcall benchmark set. For both the regions, better precision and F-measure were observed when GATK calls were made using the modified reference (as shown in **Figure 3D**). The detailed results from the evaluation tool are given in **Supplement Table 11.**

Next, we assessed the performance improvement on RNA sequencing data for the HG002 B-Lymphocyte cell type. Here we used STAR^21^ aligner within FixItFelix and subsequently GATK for variant calling. First, we assessed the coverage/expression changes across the genes (see **Supplement Table 12** for details). For falsely duplicated regions, we observed a higher coverage (2.22x more coverage on an average) for all exon regions when the modified reference was used. For falsely collapsed regions, we didn’t observe reads across the genes *FANCD2, KCNJ12* and *KCNJ18* i.e., maybe the genes are not expressed. Only regions around *DUSP2* and *MAP2K3* showed mapping reads. As expected the decoys reduced the coverage over the initially collapsed regions. Whereas for the remaining regions, the coverages were exactly the same for both references.

The evaluations of variant calling were performed by comparing GATK variant calls to draft benchmark dataset (generated by running dipcall with modified GRCh38 reference). For duplicated regions, the RNA-seq F-measure increased from 0.148 (with original reference) to 0.634 when modified reference was used. We observed a significant increase in recall (0.08 to 0.52) in those duplicated regions but also a slight decrease in precision (from 1.0 to 0.81). Across the collapsed regions we only were able to assess six SNV which showed the same performance for both the references (recall: 0.5 and precision: 1.0). See **Supplement Table 12** for details.

### The modified reference improves variant detection across ancestries

So far, we have shown a clear improvement of variant calling for HG002, an individual from European ancestry. Nevertheless, to ensure these results apply beyond the European HG002, we extended our benchmark using dipcall to eight additional individuals from the T2T Diversity Panel (see **Methods**). We first generated the draft benchmark sets for these samples by calling variants with dipcall by using hifiasm assemblies of their paternal and maternal haplotypes. This collection includes datasets for four African (HG03098, HG02055, HG02723, and HG02145), two American (HG01109 and HG01243) and two Asian (HG02080 and HG03492) samples. For the four African samples, we observed a significant improvement of F-measure for all false-duplication regions. We also observed similar improvements for the two American and two Asian samples. The whisker plots of **Figure 4A** show the F-measures for all eight samples. Overall the F-measures are significantly higher for all samples (Kolmogorov-Smirnov test *p*-value < 2.2e-16) and all duplicated regions when the modified reference is used. The regions covering the genes *CRYAA/CBS/U2AF1, ICOSLG, SIK1, SMIM11A/KCNE1* and *TRPM3* are present show nearly perfect F-measure of 1.0 for most of the samples. The F-measure comparison for SNVs of all regions of individual samples are shown in **Supplement Table 13**.

**Figure 4:**
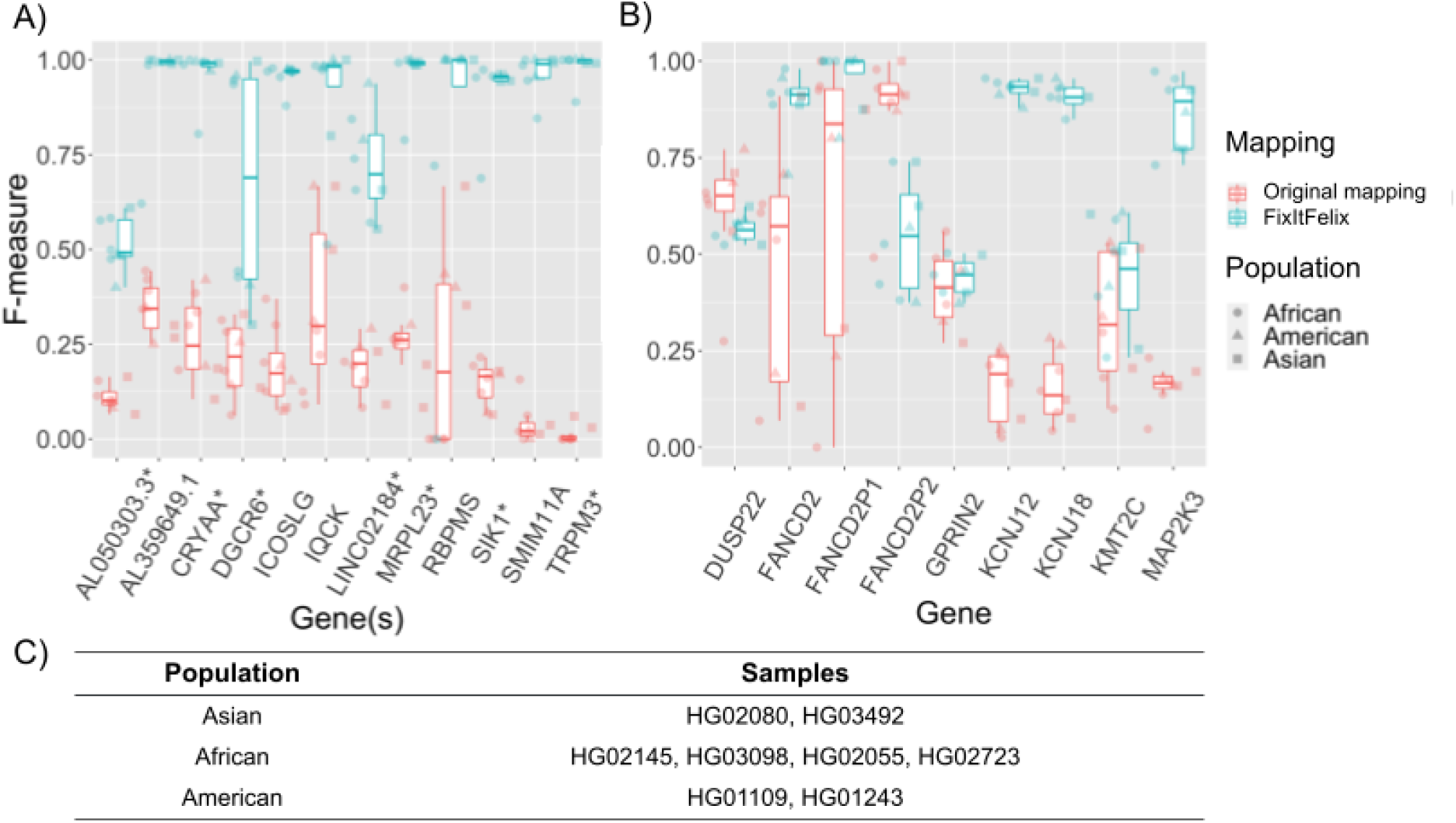
F-measures benchmarking for 8 pan-genome samples. SNV benchmarking of both **A)** duplicated and **B)** collapsed regions using original GRCh38 reference and modified reference. The x-axis is labeled with gene names that are impacted due to duplication/collapsed events. For some of the duplicated regions, more than one genes are impacted and gene names with * (star) subscripts represent several other genes including the labeled gene name. The red and green box plots correspond to original and modified references respectively. **C)** The eight pan genome samples chosen from 1KGP consists of African, American and Asian samples.

We observed a decrease in recall scores for the collapsed regions that contain *DUSP22* gene and *FANCD2P2* pseudogene. This could be due to the fact that some reads that had correctly aligned to these regions are now aligning to the corresponding decoy sequences or have too low mapping quality. However, the variant precision improved, showing the modified reference’s ability to exclude false variants that were initially called when the original GRCh38 reference was used. The above pattern was observed for all African, American and Asian samples. The collapsed regions that contain genes such as *FANCD2, KCNJ12, KCNJ18* and *MAP2K3* showed a significant improvement over F-measures (Wilcoxon Rank Sum test: *FANCD2 p*-value = 0.003824, *KCNJ12 p*-value = 0.0001554, *KCNJ18 p*-value = 0.0001554 and *MAP2K3 p*-value = 0.0001554, **Figure 4B**). This was consistent among all 8 samples that we used in our analysis. For half of the regions, the improvement was significant with an 80% increase in F-measures when modified reference was used. **Supplement Table 14** contains the F-measure comparison of all regions for individual samples. Thus, the improved accuracy across different samples and different populations demonstrates that the errors are not specific to the HG002 sample.

### New realignment allows to scale to thousands of human genomes

Given the improvements across the whole genomes for multiple ancestries, it is clear that FixItFelix, together with the modified version of GRCh38, improves variant calling and mapping in multiple important regions of the human genome. We next applied FixItFelix across 4,174 publicly available samples to measure the benefits for joint calling and obtain more insights into these genes. These samples include high coverage (target depth of 30x) sequencing of 3,199 individuals from the 1KGP^16^, 828 individuals from the Human Genome Diversity Project (HGDP)^22^, and 147 quality control sequences from the Trans-Omics for Precision Medicine program (TOPMed)^23^.

For mappings to both the original and modified reference, we calculated the mean mapping quality within each of the falsely duplicated and collapsed regions for each of the 4,174 samples. Mapping qualities for the falsely duplicated regions were consistently improved by using the modified reference (see **Supplementary Figure S1**). For the collapsed regions, the mapping quality either decreased or remained roughly the same when using the modified reference (see **Supplementary Figure S2**). These results are consistent with the evaluations of mapping quality described in previous sections.

Since falsely duplicated sequences would have mappings spread across two sequences (**Figure 1A**), it is expected that the read depths for these regions would be decreased. Likewise, reads from two different sequences would be mapped onto the same collapsed sequence, resulting in an increase in read depth for such regions. In order to assess whether this was happening in the falsely duplicated and collapsed regions that we identified, we calculated the mean depth for each variant within both call sets and compared the affected regions to the rest of the genome (see **Figure 5A**). With the original reference, the mean read depth for the collapsed regions (51.5) was approximately 1.5 times higher than the mean read depth of the unaffected regions (33.7). Similarly, the mean read depth for the falsely duplicated regions (17.7) was nearly half that of the unaffected regions. With the modified reference, these deviations in mean read depths for the collapsed and duplicated regions receded to 38.4 and 34.5 respectively.

**Figure 5:**
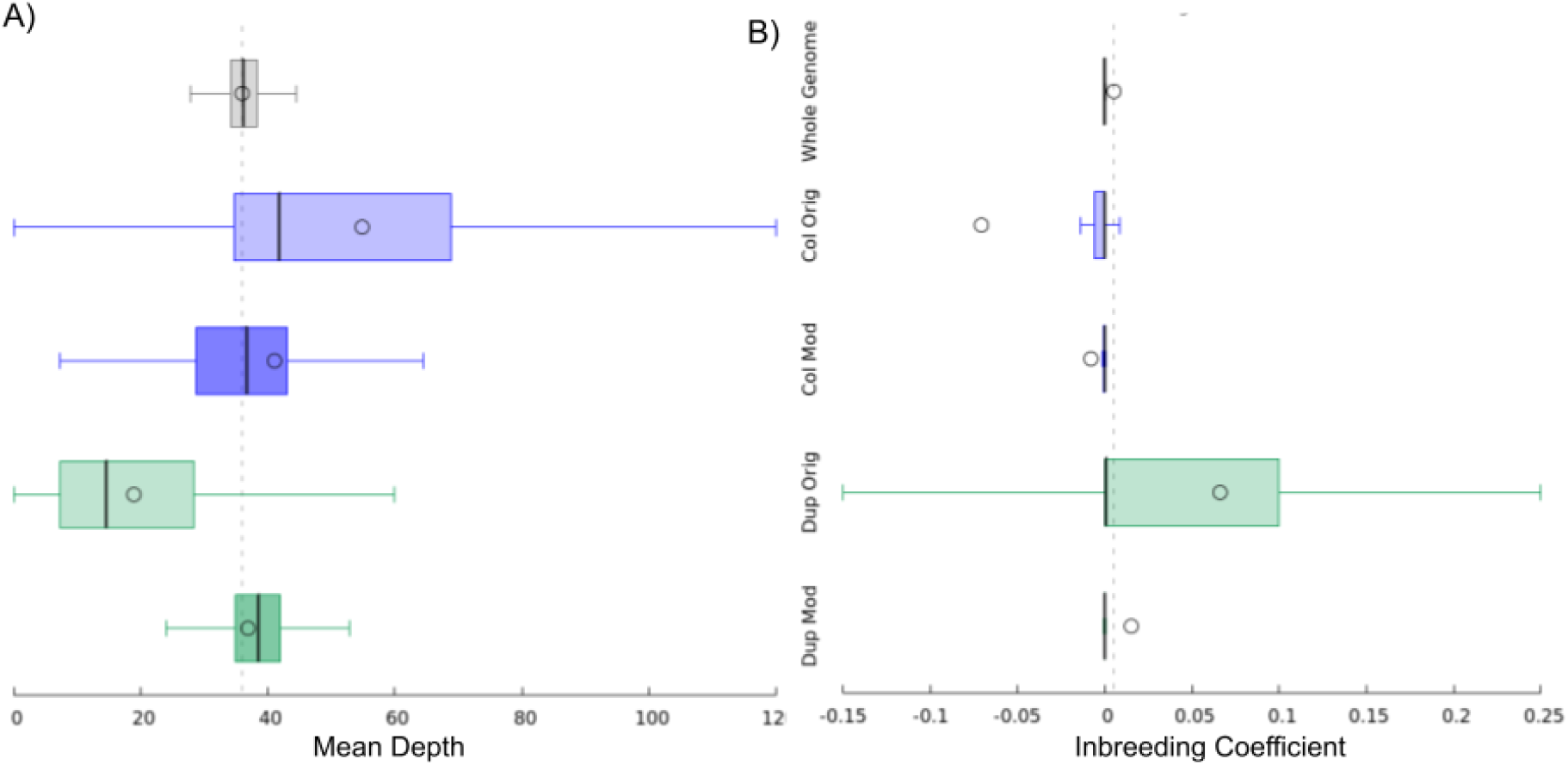
Mean read depth and inbreeding coefficient distributions. Distributions of **A)** mean read depth and **B)** inbreeding coefficient of variants for the whole genome (excluding duplicated and collapsed regions), collapsed regions using the original reference, collapsed regions using the modified reference, duplicated regions using the original reference, and duplicated regions using the modified reference. Distribution means are indicated with circles, whiskers denote min/max within -/+ 1.5 times interquartile range, and box lines denote quartile boundaries.

These types of mapping errors can often lead to artificial departure from Hardy-Weinberg equilibrium^24^, and we saw this play out in our experiment. In variant calls generated from the original mapping, we observed elevated rates of variants with heterozygous deficiency (indicated by positive inbreeding coefficients) within the falsely duplicated regions, likely caused by reads mapping to the false copy of the sequence when they have a variant that matches the false copy, which also results in lower coverage. Conversely, we observed excessive heterozygosity (indicated by negative inbreeding coefficients) within the collapsed regions, likely caused by paralogous sequence variants (PSVs) in reads mismapping to each region from the missing paralogous region^11^, also resulting in higher coverage (see **Figure 5B** and **Supplementary Figure S3**). These departures from Hardy-Weinberg equilibrium were improved for both the falsely duplicated and collapsed regions when using the modified reference.

### GRCh38 errors impact gene expression quantification and lead to artifactual cis-eQTLs

We hypothesized that errors in the original GRCh38 reference impact analyses beyond variant calling. To explore this further, we mapped 449 1KGP lymphoblastoid cell line RNA-seq datasets ^25^ (see **Supplement Table 15**) to the modified and the original reference, and compared gene read counts between the references (**Figure 6A**). As expected, when using the modified reference the number of reads mapping to (unmasked) duplicate genes generally increased. These reads commonly multi-mapped or mapped to the false paralog in the original reference (**Supplementary Figure S4**); frequently the expression of the two paralogs is negatively correlated (**Figure 6B**). This suggests that in the case of falsely duplicated regions, the false paralogs may compete with the true gene for RNA-seq reads during read mapping. Changes within collapsed regions were more restricted, with only two neighboring genes (*DUSP22* and *ENSG00000287265)* in collapsed regions showing both substantial changes in gene counts and mean counts per million reads (CPM) > 0.1 in either reference. As expression of these genes may vary across tissues, the changes observed for any one gene may vary as well.

**Figure 6:**
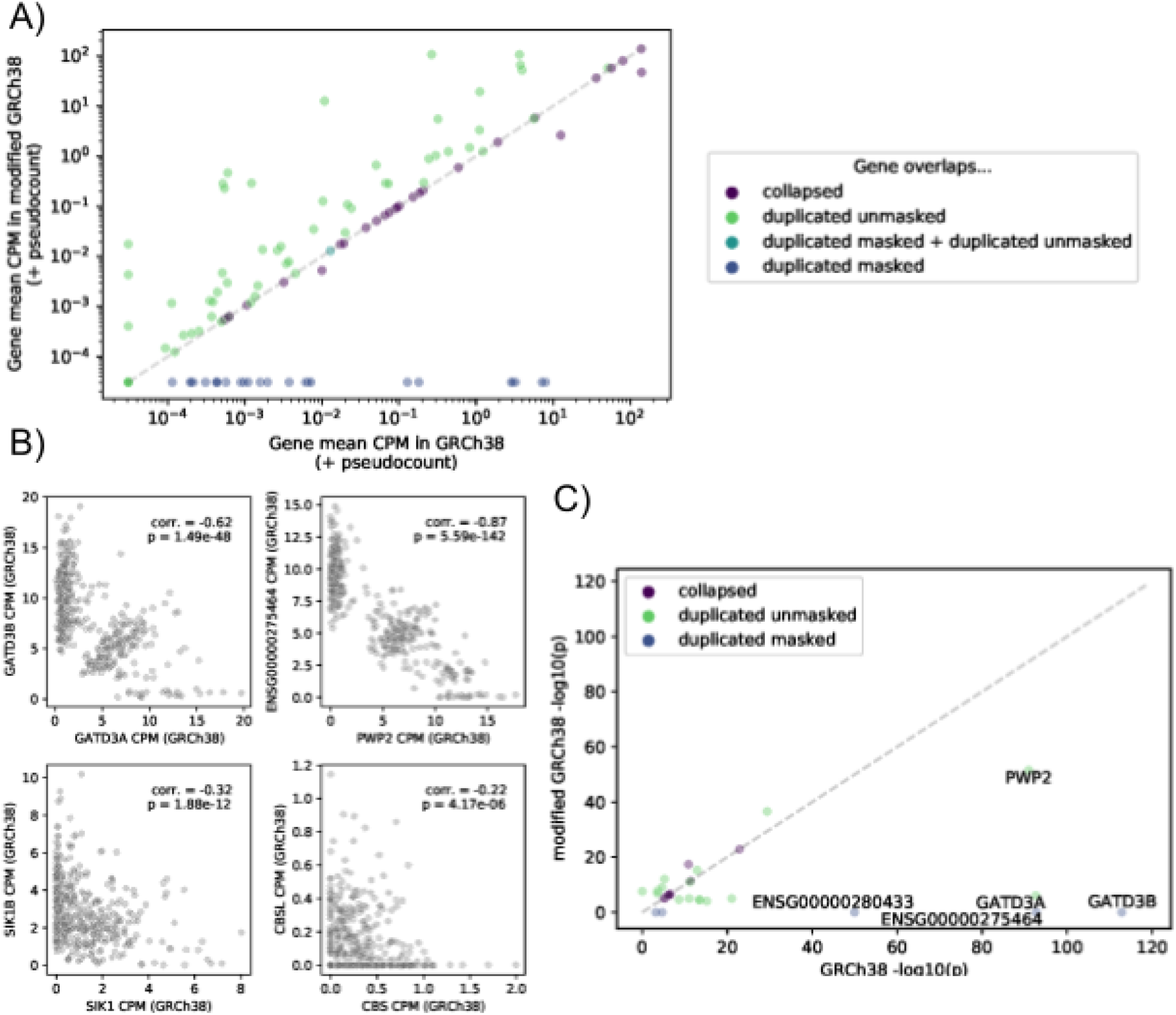
RNA-Seq eQTL analysis. **A)** Mean counts per million read pairs (CPM) across RNA-seq samples for each gene overlapping collapsed, duplicated masked, or duplicated unmasked regions, using GRCh38 or the modified GRCh38. **B)** Per-sample CPM of a true gene (x-axis) and it’s false paralog (y-axis) for four selected pairs, displaying significant negative correlation between paralogs. **C)** Comparison of cis-eQTL p-values for genetic variants most significantly associated with each gene’s expression in GRCh38 or the modified GRCh38. Only genes that are an eGenes in at least one of the references and that are within collapsed, duplicated masked, or duplicated unmasked regions are shown; if a gene was not included in the eQTL scan for one of the references (because the expression was too low), the p-value was set to 1 for the purposes of this panel. The 5 plotted genes showing the most extreme change in top cis-eQTL p-value between the references are labeled.

To demonstrate the impact of these changes on downstream analyses, we used gene counts from the modified and original reference, along with the corresponding genotypes, to perform a *cis* expression quantitative trait locus (*cis*-eQTL) analysis using these samples. We identified 10,450 genes with at least one significant eQTL signal (eGenes; 5% FDR) in at least one of the references (10,437 eGenes in GRCh38; 10,417 eGenes in modified GRCh38). Five genes in duplicated masked regions (*ENSG00000275464, ENSG00000277117, ENSG00000280433, GATD3B, SIK1B)* were significant eGenes in GRCh38. We compared the p-values for the variants most significantly associated with each gene’s expression in either reference (**Figure 6C**; **Supplementary Figure S5**; **Supplement Table 16**). As expected, genes in unaffected regions showed little change between the references. Across all genes that were an eGene in at least one of the references, 22 genes showed a change in nominal p-value (for association with the most strongly associated variant) between the references exceeding 1 order of magnitude, with only one of these (*TRAPPC10)* located outside of affected regions.

Several eQTL signals in the original reference weakened considerably in the modified reference. The *PWP2* and *GATD3A* GRCh38 cis-eQTL signals represent particularly striking examples. To explore the reason for this change, we examined the top variant–gene expression associations for GRCh38 eGenes showing a weaker eQTL nominal *p*-value in the modified reference than in GRCh38 (at least one order of magnitude smaller). For each of these genes, the RNA-seq sample genotypes of the top associated GRCh38 variant (eVariant) were always identical between the references (excluding genes masked and therefore untested in the modified GRCh38), suggesting that the decrease in eQTL signal strength was due to changes in gene expression quantification rather than genotype calling. Interestingly, for these genes we found that the change in gene read counts between the two references was often clearly associated with individuals’ genotypes, suggesting that the ability to map a read to the gene in the original reference was dependent upon genotype and the original eQTL signal was likely artifactual (**Supplementary Figure S6 & S7**). Consistent with the hypothesis that paralogs between the duplicated regions compete for RNA-seq reads in the original reference and that a reads preference for one or the other is correlated with genotype, the eVariants for the strong signals that weakened in the modified reference frequently correlated with the expression of the (often distant) paralogs, with the opposite direction of effect (**Supplementary Figure S8**). This hypothesis is also supported by the higher than expected fraction of homozygous variants in false duplications in the Hardy-Weinberg equilibrium analysis above. We therefore urge caution when interpreting gene expression analysis results for genes in affected GRCh38 results; for example, the top variants associated with *PWP2* and *GATD3A* gene expression in GRCh38 in our analysis (associations that weaken considerably when running the analysis in modified GRCh38) are also the top eVariants for these genes in one or more GTEx tissues^17^.

## Discussion

The errors due to falsely duplicated and collapsed events in the GRCh38 reference (GRCh38.p13) have greatly impacted the variant calls in many genes, including medically relevant genes (**Figure 1**). Therefore, previous studies on these impacted genes using GRCh38 might contain some erroneous information if variant calls were used for any analysis (e.g. gnomAD, GTEx, etc.) (e.g. **Figure 1D**). In this work, we focused on fixing these erroneous regions in GRCh38 (p13 release) reference by masking the extra copy of duplicated regions and adding decoys for falsely collapsed regions, thus improving among the recent release GRCh38.p14. To circumvent the need to remap to the entire genome or even larger cohorts we developed an efficient methodology FixItFelix to correct existing bam files with minutes of compute. Thus allowing for a rapid improvement across these genes. We show that the modified reference improves SNV calling for long and short reads of the whole genome, whole exome data as well as for RNA seq data. FixItFelix further can be applied also for future corrections as it is not implemented specifically for the GRCh38 genome.

As described in a recent publication^13^, we identified multiple genes that were accidentally duplicated leading to a low mapping quality and thus missing of variants along medically relevant genes and other regions of the genome. Furthermore, we collaborated with the T2T consortium to further identify multiple genes that were collapsed, enabling the identification of multiple haplotypes across genes in these regions^11^. In this study, we not only showed a solution to the two scenarios, but further tested if these are also true among the human population. We studied these artifacts across multiple individuals for whole genome and RNA sequencing.

For whole genome sequencing, we show that our approach improves variant calls across all falsely duplicated regions, and that it clearly improves some collapsed regions but others are more complex. We provide a new GRCh38 reference fasta that masks the extra copy of all false duplications and includes new decoys associated with the genes *KCNJ12, KCNJ18, KMT2C*, and *MAP2K3.* We provide an additional fasta with decoys for *FANCD2, EMC2, FANCD2P1, GPRIN2*, and *DUSP22* (see **Data availability**), which we found reduces false positives but can also substantially increase false negatives in *FANCD2P1, GPRIN2*, and *DUSP22* due to reduced mapping quality and possible structural errors in GRCh38 or structural variation.

For RNA sequencing, the evaluation remains challenging as the genes and therefore the variants that we utilized to identify true or false positives are not always expressed. Nevertheless, the coverage overall has shown benefits. Furthermore, we show that a subset of falsely-duplicated genes and genes in collapsed regions show substantial changes in estimated expression when quantifying expression with GRCh38 or the modified GRCh38, and cis-eQTL results uncovered evidence of genotype-associated mapping differences between the references that may lead to false eQTL signals when using the original GRCh38. We urge caution when interpreting the results of GRCh38-based gene expression related analyses for genes overlapping these duplicated or collapsed regions.

Ongoing efforts from T2T and other consortia are producing genomes that resolve these regions, e.g. T2T-CHM13. While we utilize this information to create the decoys that we introduced, we are also aware of all the annotational resources that are built around GRCh38 (e.g. Gnomad) to rank variants for certain phenotypes. Furthermore, millions of exons and genome sequencing datasets have been or are currently analyzed on GRCh38 (All of Us, TOPMed, etc). Thus, the re-analysis by using T2T-CHM13 on these large consortia data may not be an efficient solution. Therefore, to overcome the computational burden for large consortia our solution is ideal as it only takes a few minutes of additional analyses to improve and correct the mapping of reads in these medically relevant regions that are either collapsed or duplicated by chance. In addition, with the creation and improvements of human assemblies the likelihood is high that we might identify further issues with the GRCh references. Here, FixItFelix can be quickly adapted to include ancestry specific sequences or true errors in GRCh38 going forward. Overall, this study not only highlights the issues with GRCh38 and shows them across multiple ancestries for different genomic assays, but further provides an efficient solution that can be applied by every cohort or study. This clearly improves the resolution and accuracy of variant calling within minutes of compute and thus enables the usage of existing GRCh38 data at scale.

## Methods

### Collapsed and duplicated regions

We created reference decoys for ten genic regions with evidence of GRCh38 collapsed duplications (*TAS2R46, MAP2K3, KCNJ18, KATNAL2, FANCD2, LPA, MUC3A, KMT2C, GPRIN2*, and *DUSP22*). First, we identified syntenic regions in T2T-CHM13 (**Supplement Table 1**) by BLAT^26^ comparisons and matching gene annotations. Next, using T2T-CHM13 annotated segmental duplications^27^, we identified all homologs, extracted their sequences and gene annotations (UCSC Genome Browser Table Browser; T2T-CHM13 v1.0), and compared them to GRCh38 using a combination of BLAT and minimap2 ^28^. Finally, syntenic duplicate regions between references were flagged as those sharing the same gene annotations and highest sequence similarity via manual curation. T2T-CHM13 sequences of duplicate regions not represented in GRCh38 were included as decoys. Further, all reads mapping to GRCh38 syntenic duplicate regions were extracted from original bams and remapped to the new modified GRCh38 reference. We chose not to include *TAS2R46*, *KATNAL2*, *LPA*, and *MUC3A* because they did not have simple decoy sequences that could be added.

### Remapping tool - FixItFelix

Our remapping tool, FixItFelix, extracts only the mappings of the regions of interest from the existing whole genome mapping BAM/CRAM file and then extracts sequences for those regions (duplicated or collapsed) and finally realigns the sequences to the modified GRCh38 reference. This is significantly faster (5~8 mins) than global whole genome mapping (may take >24 hours). Following are the detailed steps of FixItFelix (see **Figure 1B**). First, the regions in the original BAM/CRAM file, i.e. mapped to GRCh38 reference genome, corresponding to input BED regions (duplicated or collapsed) are extracted using samtools ^29^ (v1.12) with “-F 2316” flag (primary alignments, not supplementary alignments, reads are not unmapped, and mate pairs are not unmapped). The extracted alignments were further filtered to make sure that we are keeping the alignments only for paired-end reads with unique read names. Then, the sequences are extracted from these alignments using the “samtools fastq*”* command. Finally, the extracted reads were mapped to the modified reference genome using BWA^30^ (v0.7.15).

### Short read mapping genome wide

For comparing the mapping quality, we first aligned the reads to entire reference genomes (original and modified GRCh38). The short reads were mapped to both original and modified reference using the bwa-mem algorithm of BWA (v0.7.15) aligner tool with minimum seed length (-K) set to 100,000,000 and all other parameters were set to its default value. The mapping qualities were evaluated using a customized script that used the samtools (v1.12). We also manually examined a few of the mapping regions of our interest using the Integrative Genomic Viewer (IGV) tool (v2.12.3).

~~~
bwa mem -K 100000000 -Y -t 8 -R
@RG\tID:0\tSM:HG002\tLB:HG002\tPU:HG002_38_nodecoy\tCN:BCM\tDT:2021-03-10T00:0 0:00-0600\tPL:Illumina GCA_000001405.15_GRCh38_no_alt_analysis_set.fasta HG002.novaseq.pcr-free.35x.R1.fastq.gz HG002.novaseq.pcr-free.35x.R2.fastq.gz
~~~

### Variant calling

To evaluate the variant calls when mapped to the original and modified reference, we used both genome wide mapping and the mappings corresponding to regions impacted by duplicated and collapsed events. We use our remapping script to extract BAM regions from both the original mapping and the mapping to the modified reference genome.

Single-sample variant calling for whole genome, regional and remapped regions was done using GATK (v3.6) Haplotypecaller^31^. Jointly genotyped call sets for 1KGP, HGDP, and TOPMed samples were generated with the TOPMed variant calling pipeline. The customized scripts for variant calling are available at our github repository (see **Code availability**).

### Long-read variant calling

We called variants (SNVs and indels) using existing long-read aligned bam file and PRINCESS (v1.0)^32^, with the “snv” option and default parameters. The “snv” option from PRINCESS calls implicitly Clair3^33^ (v3.0.1.11); for the specific regions of interest (used the ‘`--bed_fn`’ option to specify the bed file).

### Evaluating variant calls

Our evaluation for HG002 dataset done using two different benchmark sets: A) GIAB challenging medically relevant gene (CMRG) benchmark set and B) dipcall benchmark set. The dipcall benchmark was chosen as CMRG covers only a small set of regions. For 1KGP samples, HG002 WES dataset and HG002 RNA-Seq dataset, we used only the dipcall benchmark set.

Using GIAB CMRG (v1.0) dataset, the benchmarking of variant calls were performed using the hap.py^34^ tool (v0.3.14) that used the high confidence region BED file of CMRG set (`-f` parameter) and duplicated/collapsed BED regions (`-T` parameter). The reference (`-r` parameter) was appropriately chosen for original mapping and mapping to modified reference.

For the eight 1KGP samples, we first run dipcall (v0.3) by taking their publicly available maternal and paternal assemblies (hifiasm tool and PacBio HIFI reads were used for assemblies). The dipcall was run with the modified reference genome and the assemblies, the output VCF file and BED file were used as benchmark set and high confidence regions for evaluation. We again used the hap.py tool for comparing the GATK VCF file with the benchmark VCF file. Our analysis on 1KGP was performed by using the dipcall benchmark set that was generated using the modified reference genome.

For WES and RNA-Seq experiments, we used HG002 dipcall results with the modified reference as the benchmark set. The bedtools (bedtools intersect) was used to extract the exon regions that overlap with duplicated/collapsed regions. The comparison of VCF files containing GATK variant calls and the truth set (dipcall benchmark) were done using the hap.py tool. For RNA-Seq, we followed a specific pipeline of GATK that was designed for RNA-Seq experiments which was different from genome wide variant calls.

The read depth and Hardy-Weinberg Equilibrium metrics used when evaluating variants in the jointly genotyped call sets were taken from VCF INFO fields that were generated with the TOPMed variant calling pipeline. Specifically, the average read depths and inbreeding coefficients for each variant were taken from the INFO fields respectively named “AVGDP” and “FIBC_P”^23^.

### Statistical testing of improvement of mapping quality and evaluated variants

We evaluated the mapping quality of improvement of FixItFelix for both duplicated and collapsed regions using the data from **Supplement Tables 3 and 4** respectively. We compared the mapping quality of both strategies using a Wilcoxon rank sum test (in R)

~~~
> wilcox.test(original.mapping.quality, fixItFelix.mapping.quality, alternative=“two.sided”)
~~~

The scores of the SNV and INDEL calls benchmark were compared using a Friedman rank sum test (in R) where we compared the F-scores by the mapping strategy (Original, Global remapping, FixItFelix) and type of event (SNP, or INDEL) for both the collapsed and masked regions (**Supplement Table S17**).

~~~
> friedman.test(F1.score ~ Mapping | Type, grch38_f1scores_supp_table_S17)
~~~

Next, we compared the F-score across 12 and nine genes for duplicated and collapsed regions respectively in eight individuals from three distinct ancestries (two from Asian, four from African and two from American ancestry). For both duplicated and collapsed regions we aggregated the F-scores for each mapping strategy (Original and FixItFelix) and performed a Kolmogorov-Smirnov test. Finally, for the case of collapsed regions, we analyzed four genes (FANCD2, KCNJ12, KCNJ18 and MAP2K3) which results looked very promising and performed a Wilcoxon test (in R) to compare the F-scores of each gene

~~~
> wilcox.test(gene.original.Fscore, gene.FixItFelix.Fscore, alternative=“two.sided”)
~~~

### Gene expression quantification

Raw RNA-seq reads were mapped to either reference using STAR^21^ (v. 2.6.1d; parameters
--outFilterMultimapNmax 20 --alignSJoverhangMin 8 --alignSJDBoverhangMin 1 --outFilterMismatchNmax 999 --outFilterMismatchNoverLmax 0.1 --alignIntronMin 20 --alignIntronMax 1000000 --alignMatesGapMax 1000000 --outFilterType BySJout --outFilterScoreMinOverLread 0.33 --outFilterMatchNminOverLread 0.33 --limitSjdbInsertNsj 1200000 --outSAMstrandField intronMotif --outFilterIntronMotifs None --alignSoftClipAtReferenceEnds Yes --quantMode TranscriptomeSAM GeneCounts --outSAMtype BAM Unsorted --outSAMunmapped Within --chimSegmentMin 15 --chimJunctionOverhangMin 15 --chimOutType Junctions WithinBAM SoftClip --chimMainSegmentMultNmax 1). STAR indices were produced using the GENCODE v. 39 GTF file ^35^(which was used for all gene expression quantification and eQTL analyses) with option `--sjdbOverhang 100`. Gene counts were quantified using RNASeQC ^36^ (v2.3.4; options --stranded rf; for use with RNASeQC the GTF was collapsed using the GTEx^17^ script (github commit 9c6a1c38b)).

To compare read feature assignments for affected genes (used for **Supplementary Figure S4**), we extracted reads that overlapped affected genes or affected regions in either reference (`samtools view -F 2048 -F 256 -L affected_regions_and_genes.bed`) and used htseq-count^37^ (v0.12.3) to assign them to features (`--stranded=yes --type=gene -a 0 --samout=out.sam`), extracting feature assignments from the XF tag in the output sam file.

### cis-eQTL analysis

We used sex, four genotype principle components (PCs), and 30 gene expression PCs as covariates for the cis-eQTL analysis. The number of gene expression PCs to include as covariates was determined by running the cis-eQTL scans using anywhere between 0 and 100 PCs (in steps of 5) and selecting the point at which the number of eGenes discovered began to saturate (**see Supplementary Figure S9**).

Gene expression values used in the cis-eQTL scan were pre-processed as follows:

1. Gene counts were filtered to include only autosomal and chrX genes
2. Genes counts were normalized using the edgeR^38^ Trimmed Mean of M-values(TMM) procedure i.e. computes normalization factors that represent sample-specific biases, as implemented in pyqtl (v0.1.8) function edger_cpm.
3. Lowly expressed genes, defined as those where < 20% of samples have a transcript per million (TPM) value of > 0.1, were dropped.
4. TMM-normalized gene expression values were inverse normal transformed.

To generate gene expression PCs to be used as covariates in the cis-eQTL scans, we performed PC analysis (PCA) on the inverse normal transformed gene expression matrix.

Genotype PCA was performed using genotypes in unaffected regions, such that the PCs for the GRCh38 and modified GRCh38 eQTL scans were identical. Genotypes for PCA were generated by filtering to common (MAF >= 1%) autosomal SNPs, followed by LD pruning using plink (`v. 1.90b; --indep-pairwise 200 100 0.1`). EIGENSOFT ^39,40^(git commit 09ed563f) was used for the PCA, computing the top 15 PCs (smartpca.perl with options `-k 15 -m 0`).

cis-eQTL scans were performed using tensorQTL^41^ (slightly modified from v1.0.6; mode = cis, with q-value lambda = 0, seed = 2021 and otherwise default parameters), testing variants within 1 Mb of the gene TSS and with in-sample MAF >= 1%.

## Data availability

Challenging medically relevant regions (CMRG) benchmark for HG002 sample (GRCh38) with high-confidence regions https://ftp-trace.ncbi.nlm.nih.gov/giab/ftp/release/AshkenazimTrio/HG002_NA24385son/CMRG_v1.00/GRCh38/SmallVariant/

GRCh38 original reference https://ftp-trace.ncbi.nlm.nih.gov/ReferenceSamples/giab/release/references/GRCh38/GCA_000001405.15_GRCh38_no_alt_analysis_set.fasta.gz

GRCh38 modified reference (used for all analysis in this study): https://bcm.box.com/s/xi95ahgzrw86pvogm7sdwl0ppn49i5dn

2nd version of Modified GRCh38 reference that excludes decoys related to *FANCD2, DUSP22* and *GPRIN2* genes https://bcm.box.com/s/qz5h36ry4cg9j15mwolzcpf1z7ha823e

HG002 HiFiasm assembly used for dipcall https://ftp-trace.ncbi.nlm.nih.gov/giab/ftp/release/AshkenazimTrio/HG002_NA24385_son/CMRG_v1.00/hifiasm-assembly/

1KGP samples: https://github.com/human-pangenomics/hpgp-data

Whole Exome Sequencing (hiseq4000,wes_agilent,50x,HG002,grch38) https://storage.googleapis.com/brain-genomics-public/research/sequencing/grch38/bam/hiseq4000/wes_agilent/50x/HG002.hiseq4000.wes-agilent.50x.dedup.grch38.bam

WES high-confidence BED regions https://www.biorxiv.org/content/biorxiv/early/2020/12/16/2020.12.15.356360/DC2/embed/media-2.gz?download=true

HG002 RNA-Seq data https://www.coriell.org/0/Sections/Search/Sample_Detail.aspx?Ref=GM24385

HG002 RNA-Seq coding regions BED https://ftp-trace.ncbi.nlm.nih.gov/ReferenceSamples/giab/release/genome-stratifications/v3.0/GRCh38/FunctionalRegions/GRCh38_refseq_cds.bed.gz

## Code availability

FixItFelix: https://github.com/srbehera/FixItFelix

Scripts: https://github.com/srbehera/GRCh38_Paper_scripts

TOPMed variant calling pipeline: https://github.com/statgen/topmed_variant_calling

Pyqtl: https://github.com/broadinstitute/pyqtl

GTEx script: https://github.com/broadinstitute/gtex-pipeline/blob/master/gene_model/collapse_annotation.py Modified tensorQTL: https://github.com/porchard/tensorqtl/commit/1822701b

## Acknowledgments

We want to thank the TOPMed IRC, as well as Severine Catreax, Fred Farrell, Justin Wagner, Alaina Shumate, and others from the T2T Consortium, for helpful discussions. Certain commercial equipment, instruments, or materials are identified to specify adequate experimental conditions or reported results. Such identification does not imply recommendation or endorsement by the National Institute of Standards and Technology, nor does it imply that the equipment, instruments, or materials identified are necessarily the best available for the purpose.

## Author contributions

F.J.S., and J.M.Z. designed and directed the study, M.Y.D., and D.C.S. worked on identifying collapsed and duplicated regions, S.B., and F.J.S. developed the remapping tool, S.B., J.L., and P.O. performed the analysis, All authors helped in the analysis, manuscript writing and provided critical feedback.

