## Supplementary Figures for "Fixing reference errors efficiently improves sequencing results"

### Supplement Figures

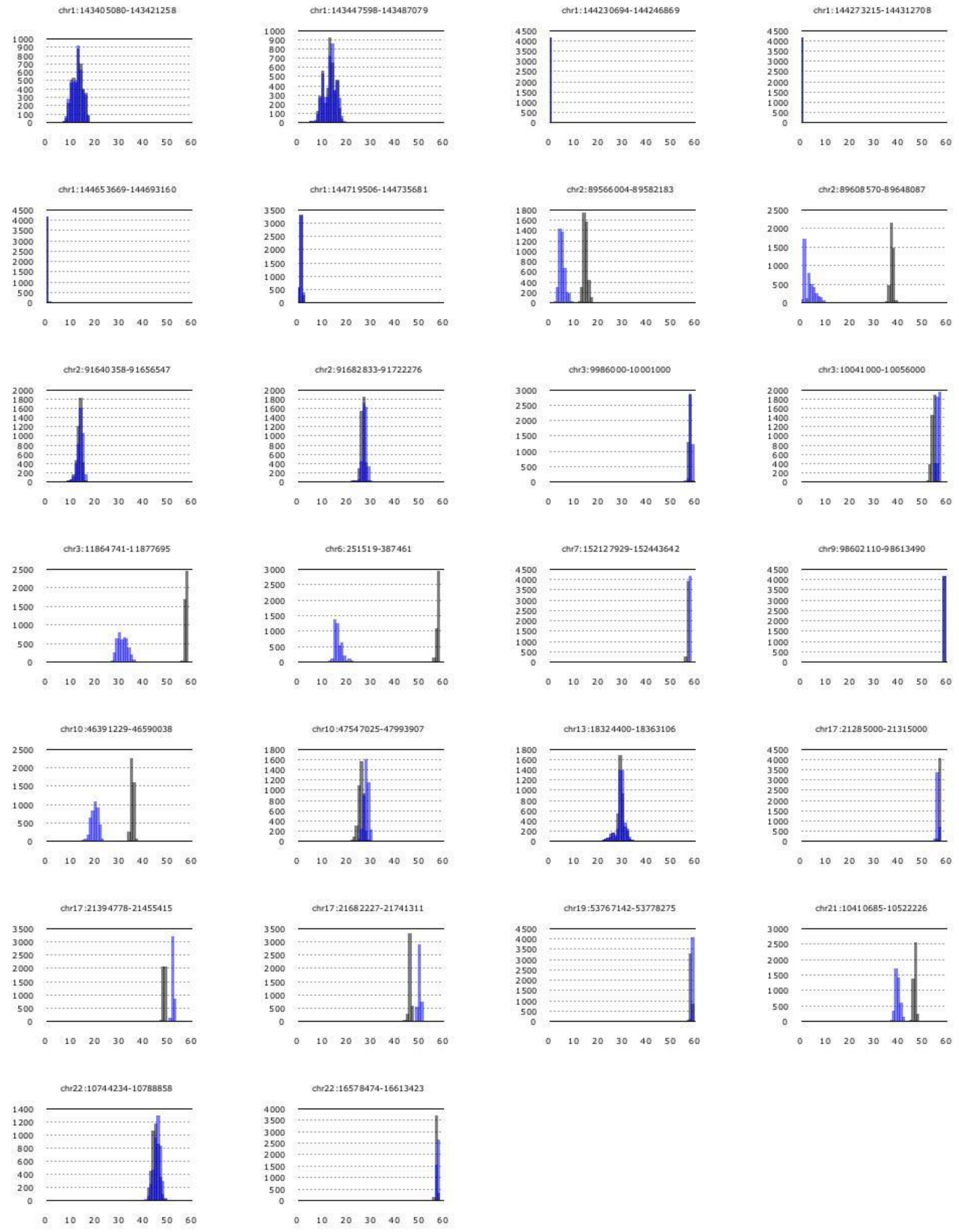


***Supplementary Figure S1: Distributions of mean mapping quality (collapsed).*** *Distributions of mean mapping quality of each sample for collapsed regions.*

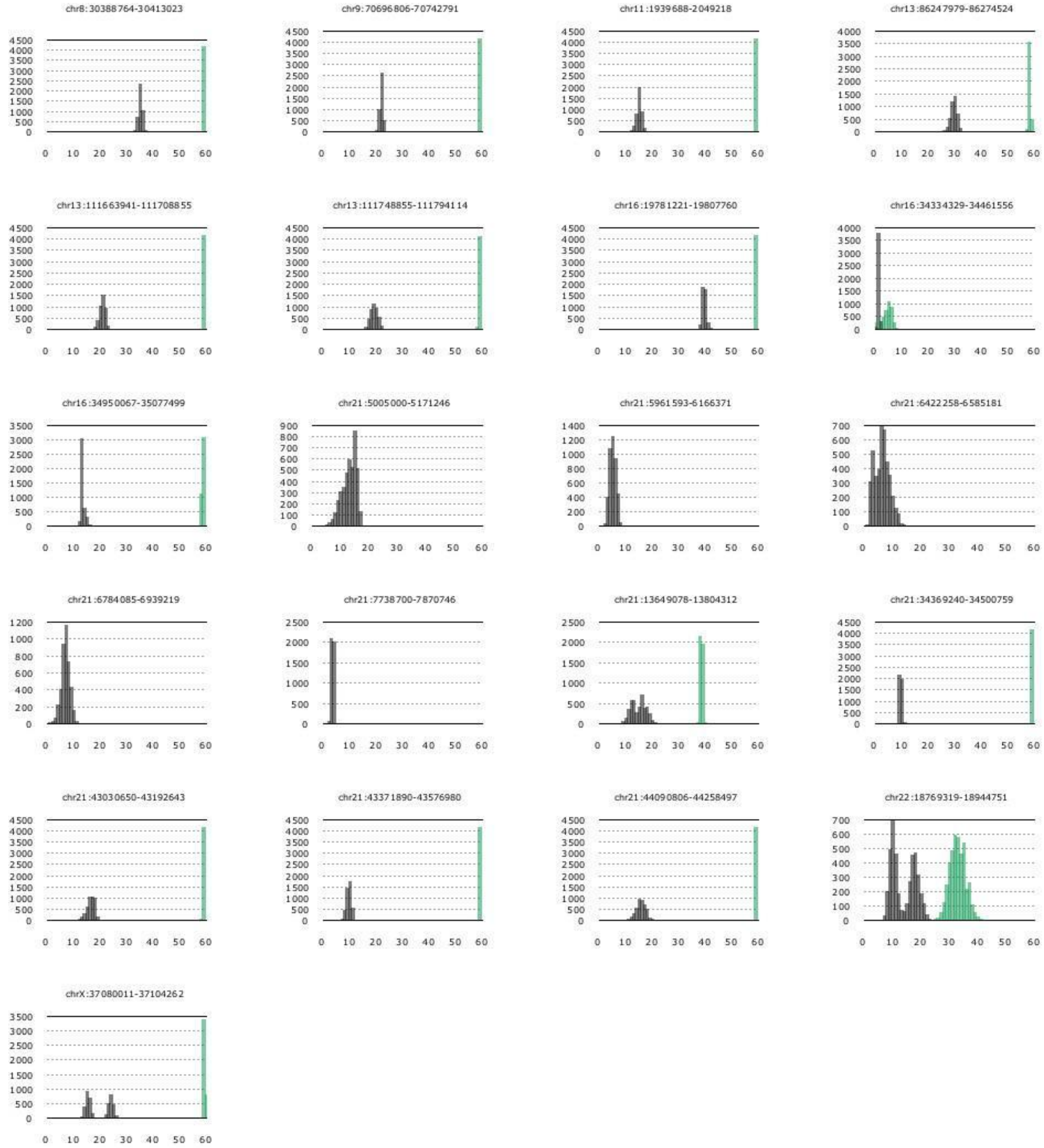


***Supplementary Figure S2: Distributions of mean mapping quality (duplicated).***
*Distributions of mean mapping quality of each sample for duplicated regions.*

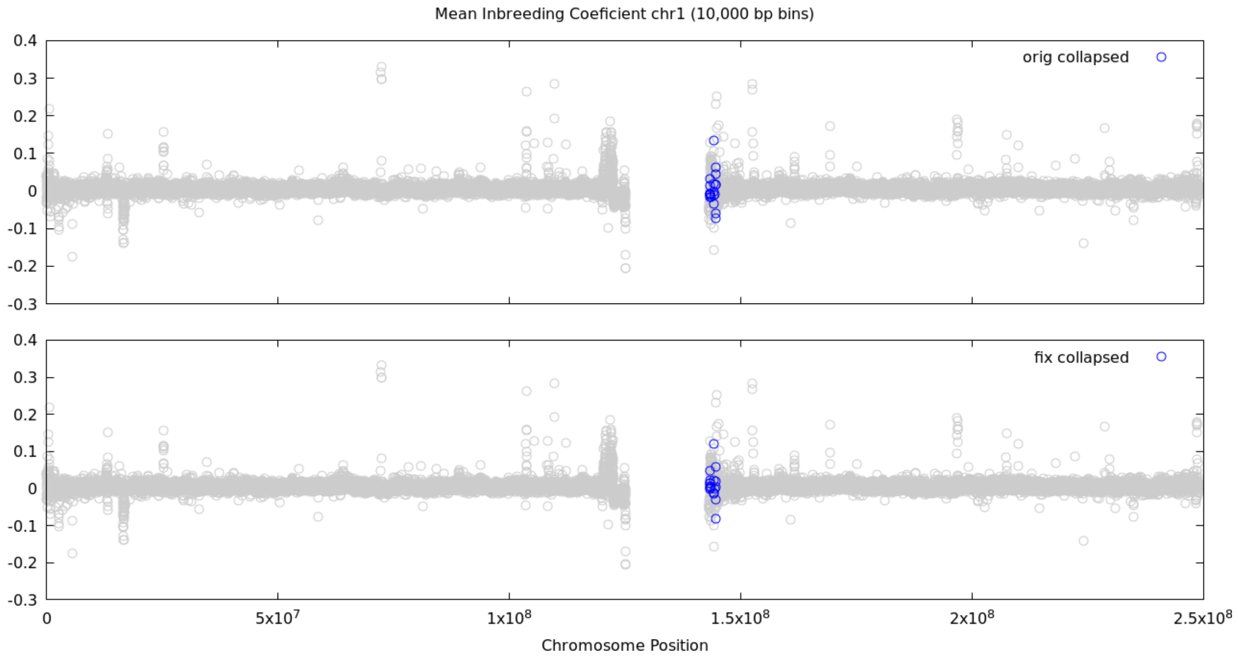


***Supplementary Figure S3.a: Binned inbreeding coefficient.*** *Mean inbreeding coefficients for 10,000 basepair bins of chromosome 1.*

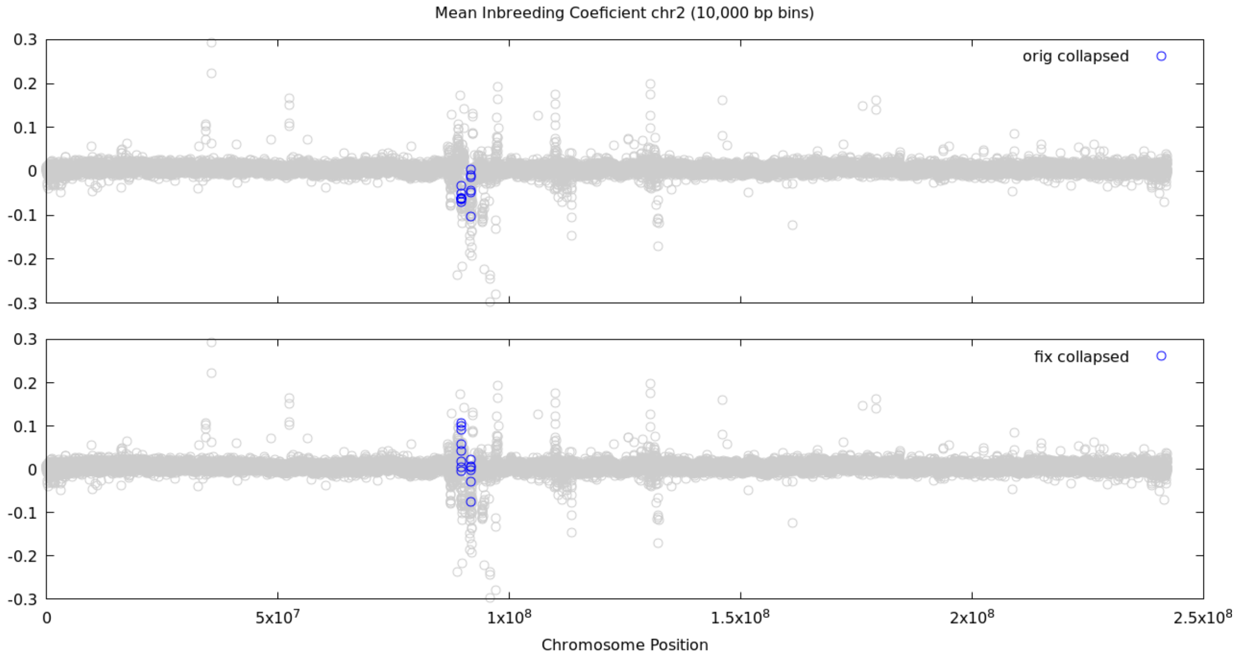


***Supplementary Figure S3.b: Binned inbreeding coefficient.*** *Mean inbreeding coefficients for 10,000 basepair bins of chromosome 2.*

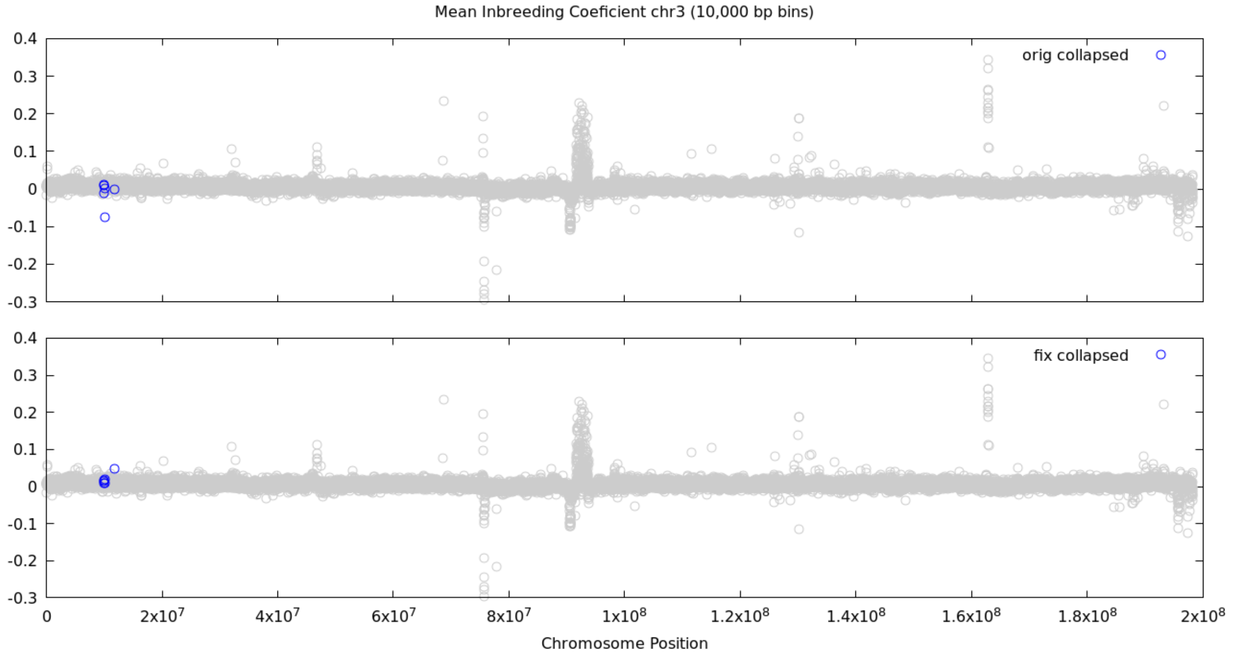


***Supplementary Figure S3.c: Binned inbreeding coefficient.*** *Mean inbreeding coefficients for 10,000 basepair bins of chromosome 3.*

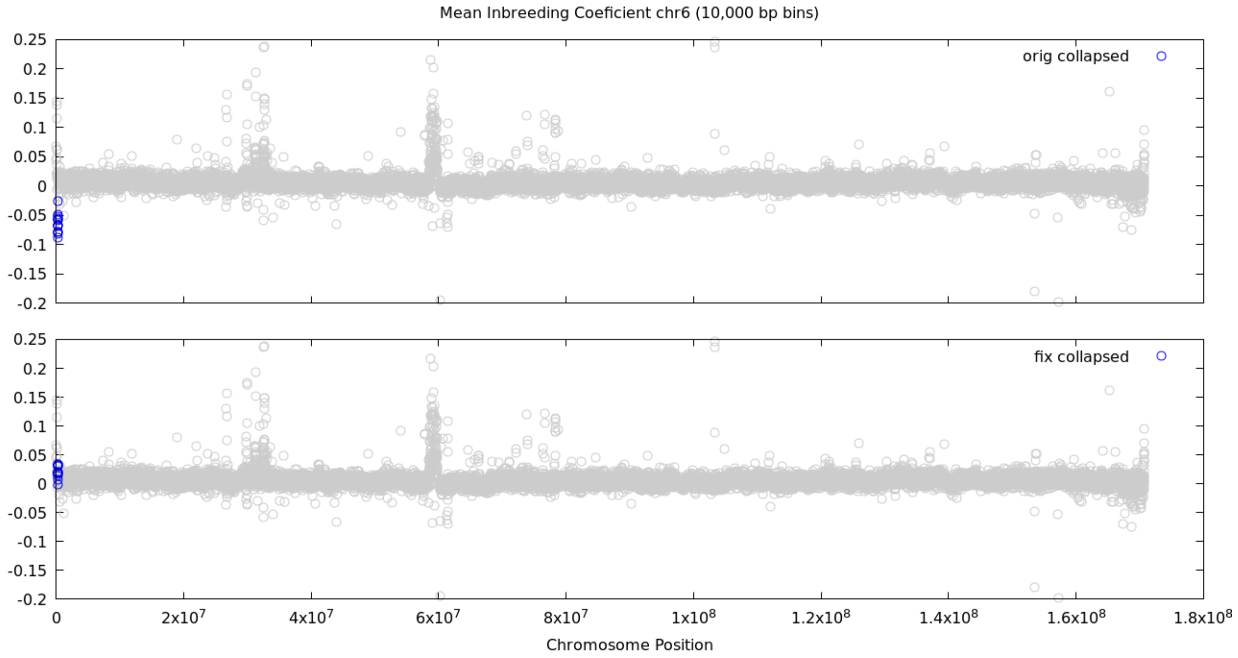


***Supplementary Figure S3.d: Binned inbreeding coefficient.*** *Mean inbreeding coefficients for 10,000 basepair bins of chromosome 6.*

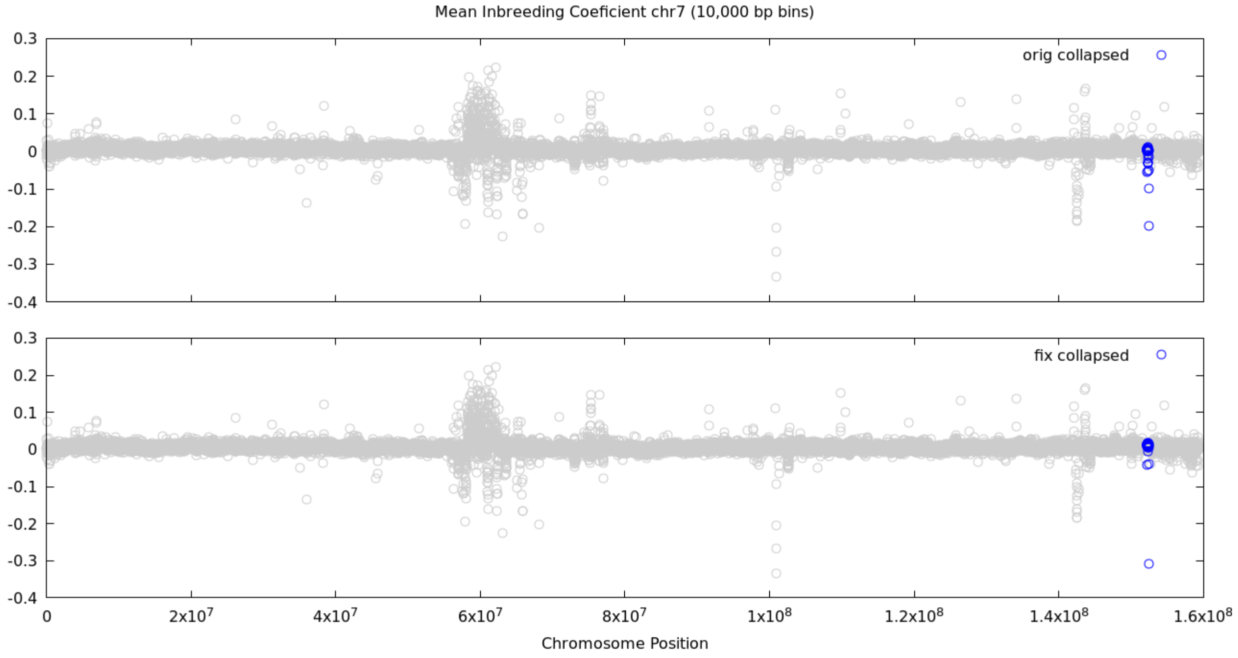


***Supplementary Figure S3.e: Binned inbreeding coefficient.*** *Mean inbreeding coefficients for 10,000 basepair bins of chromosome 7.*

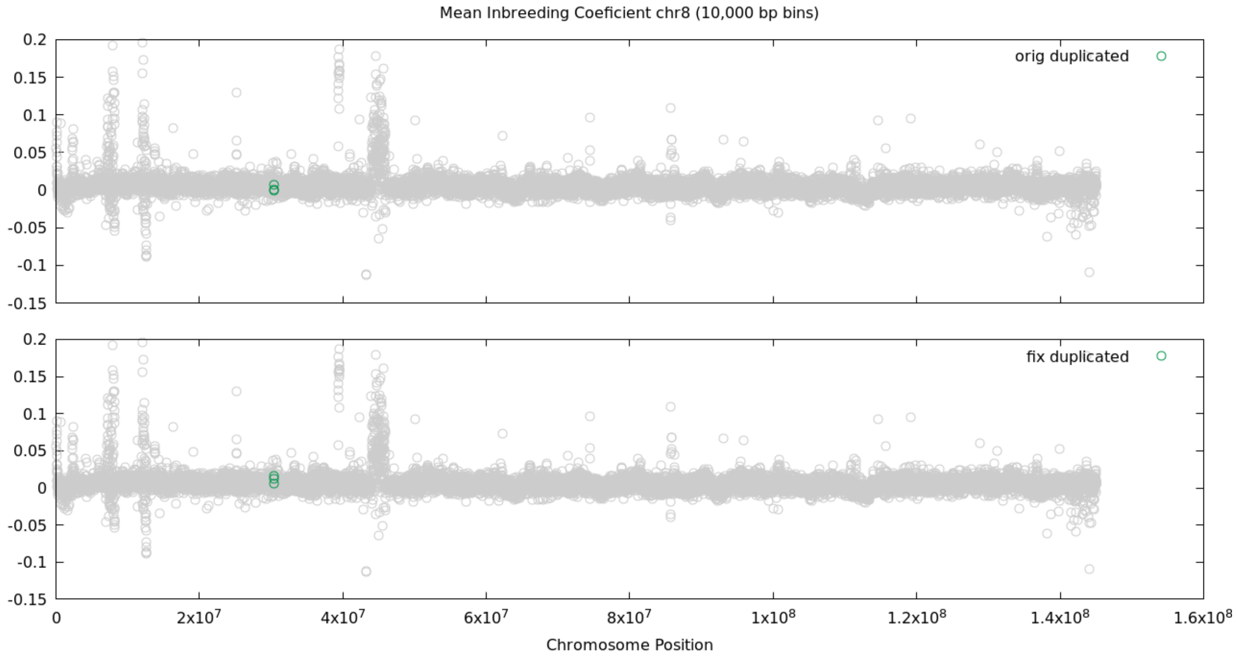


***Supplementary Figure S3.f: Binned inbreeding coefficient.*** *Mean inbreeding coefficients for 10,000 basepair bins of chromosome 8.*

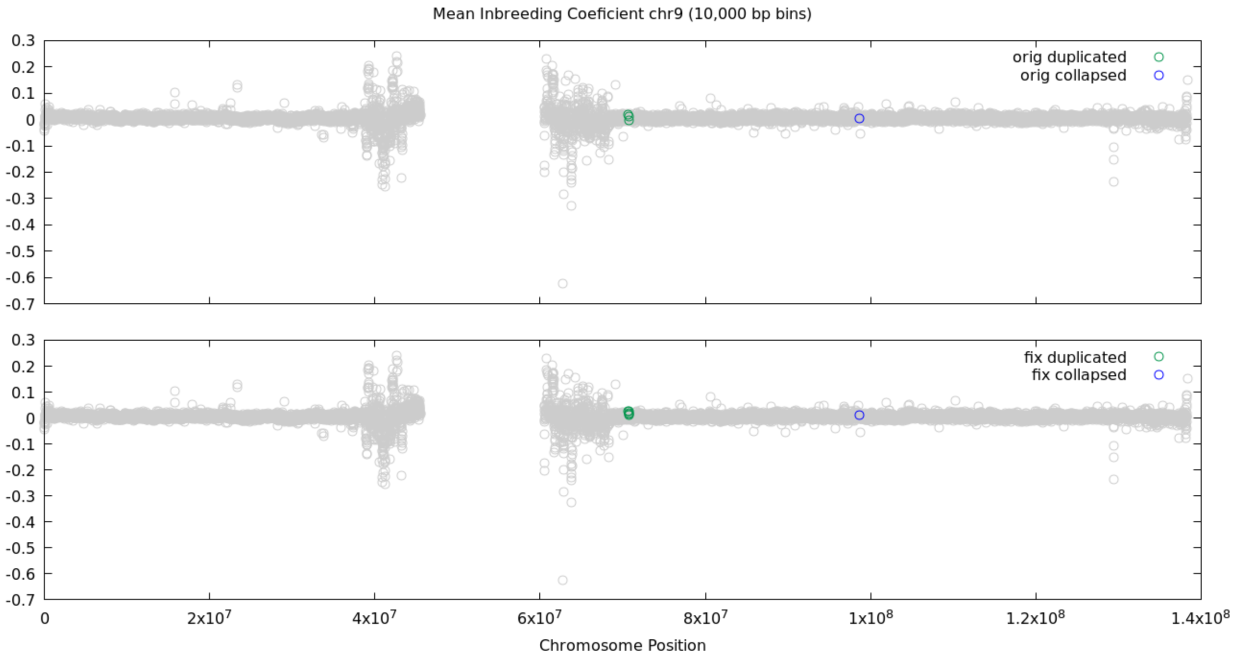


***Supplementary Figure S3.g: Binned inbreeding coefficient.*** *Mean inbreeding coefficients for 10,000 basepair bins of chromosome 9.*

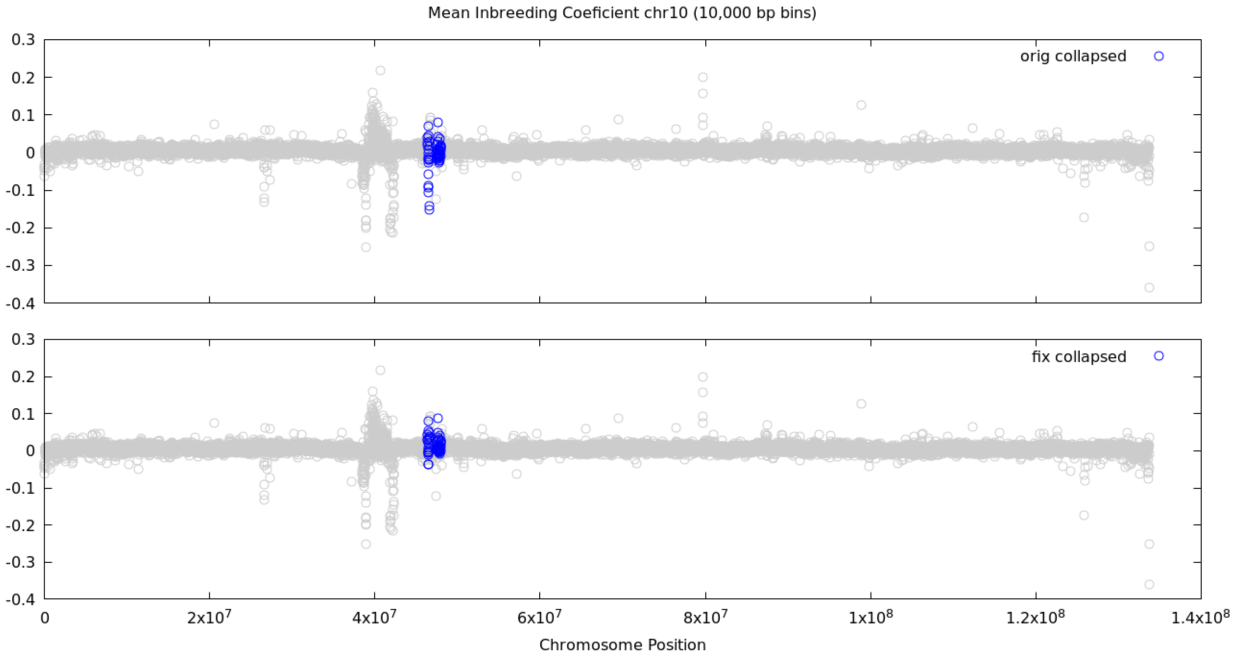


***Supplementary Figure S3.h: Binned inbreeding coefficient.*** *Mean inbreeding coefficients for 10,000 basepair bins of chromosome 10.*

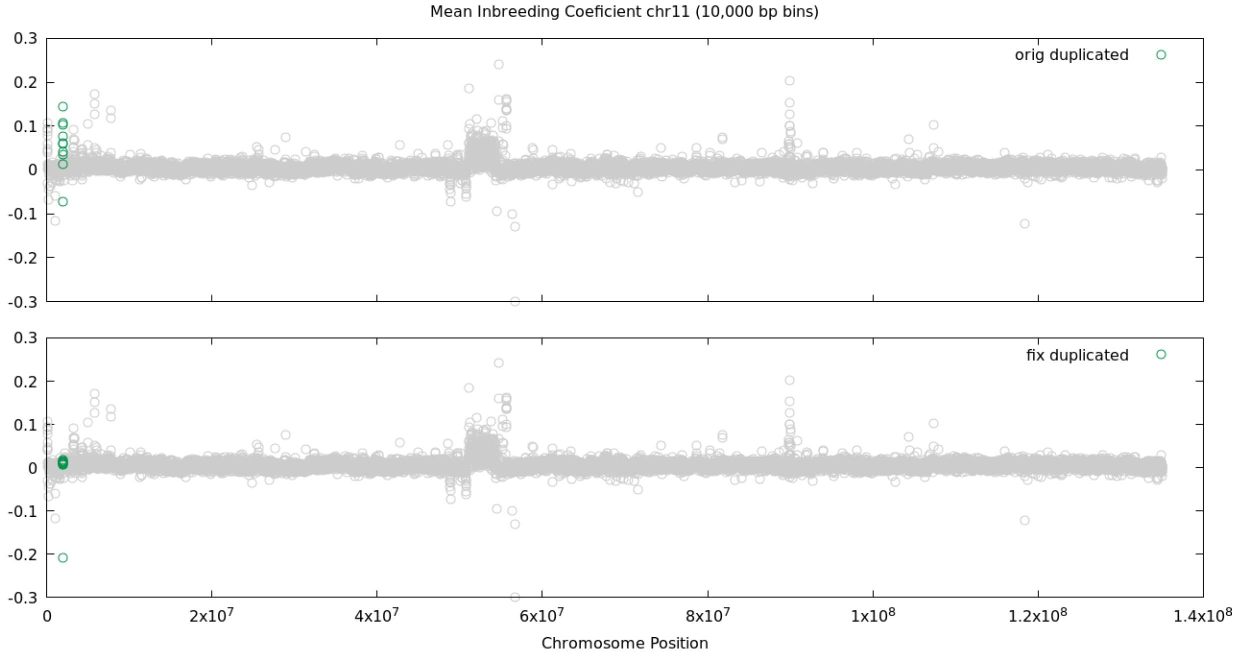


***Supplementary Figure S3.i: Binned inbreeding coefficient.*** *Mean inbreeding coefficients for 10,000 basepair bins of chromosome 11.*

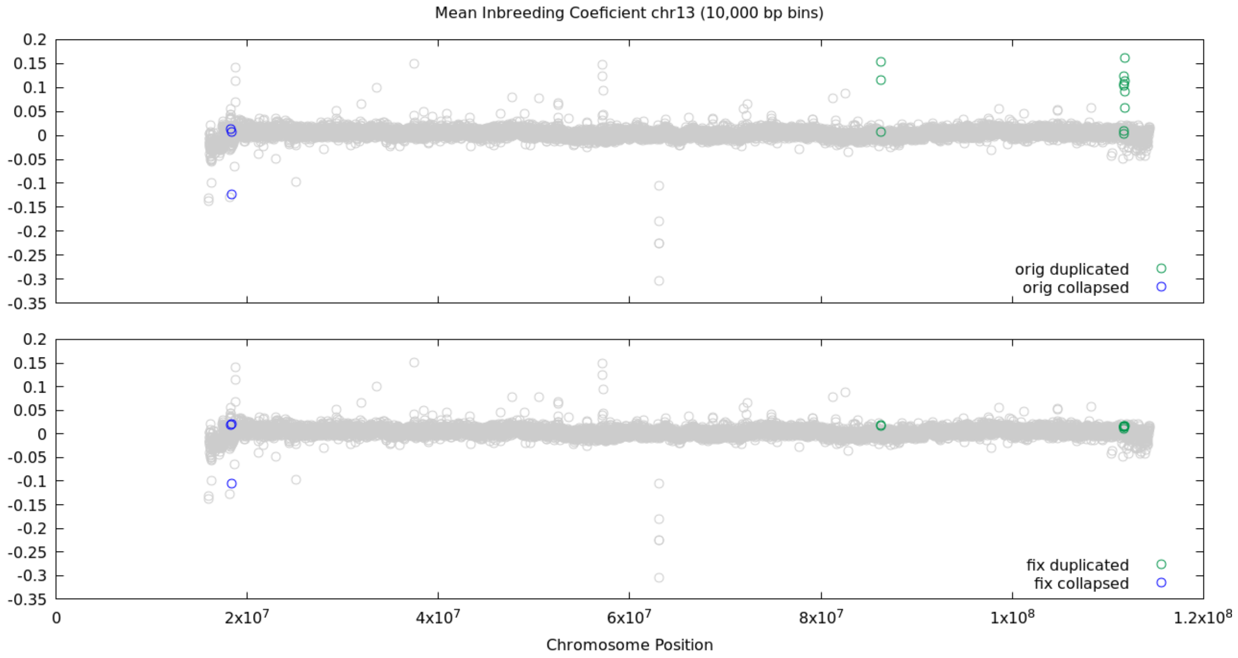


***Supplementary Figure S3.j: Binned inbreeding coefficient.*** *Mean inbreeding coefficients for 10,000 basepair bins of chromosome 13.*

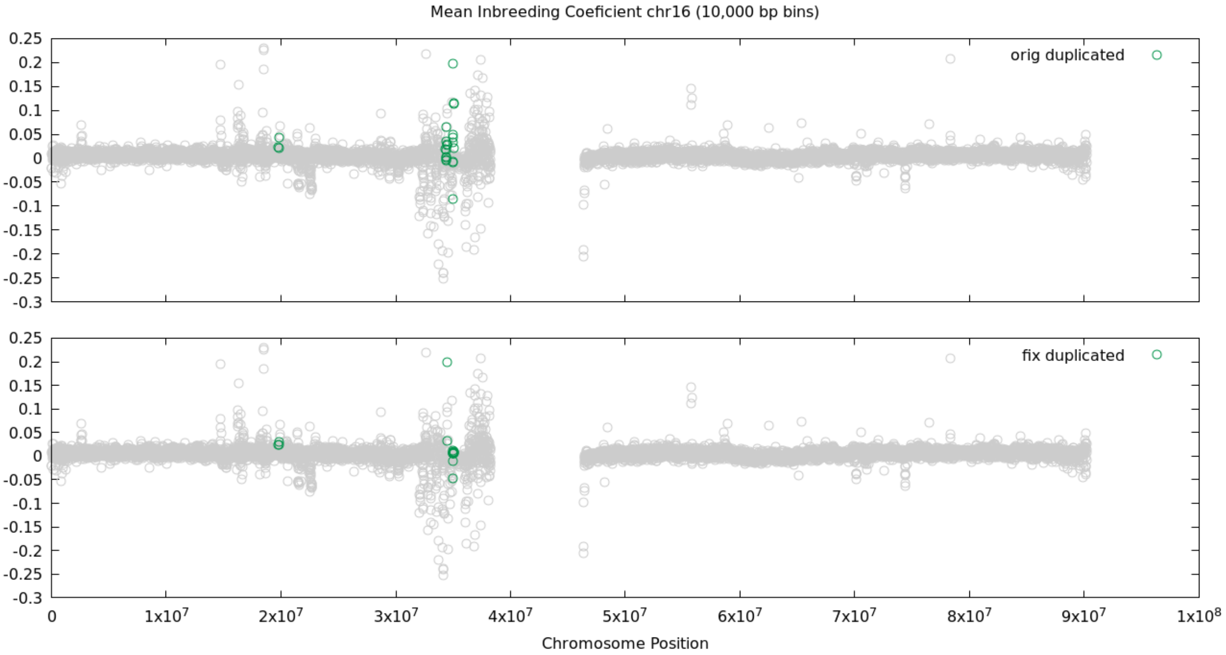


***Supplementary Figure S3.k: Binned inbreeding coefficient.*** *Mean inbreeding coefficients for 10,000 basepair bins of chromosome 16.*

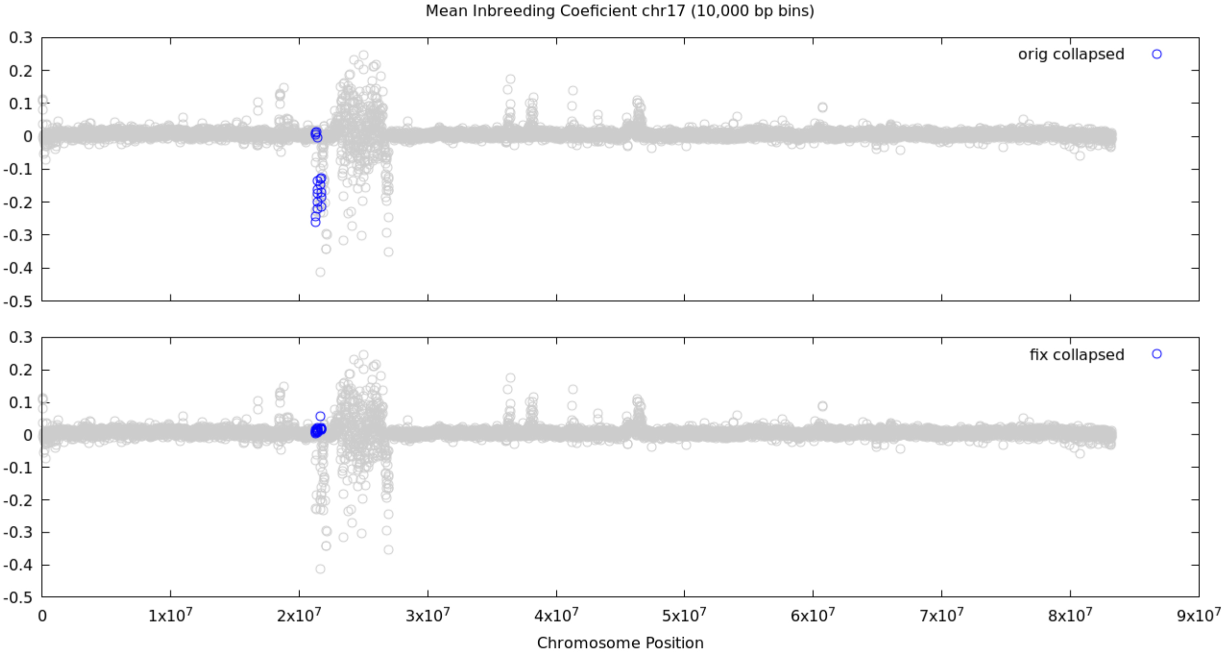


***Supplementary Figure S3.l: Binned inbreeding coefficient.*** *Mean inbreeding coefficients for 10,000 basepair bins of chromosome 17.*

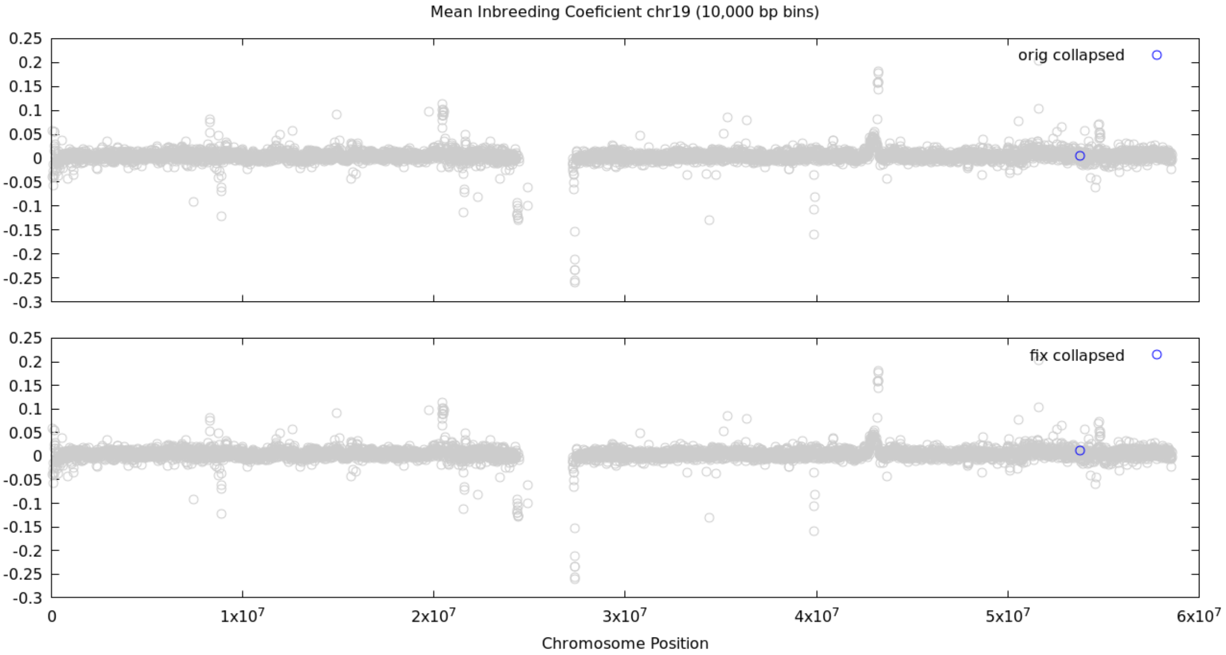


***Supplementary Figure S3.m: Binned inbreeding coefficient.*** *Mean inbreeding coefficients for 10,000 basepair bins of chromosome 19.*

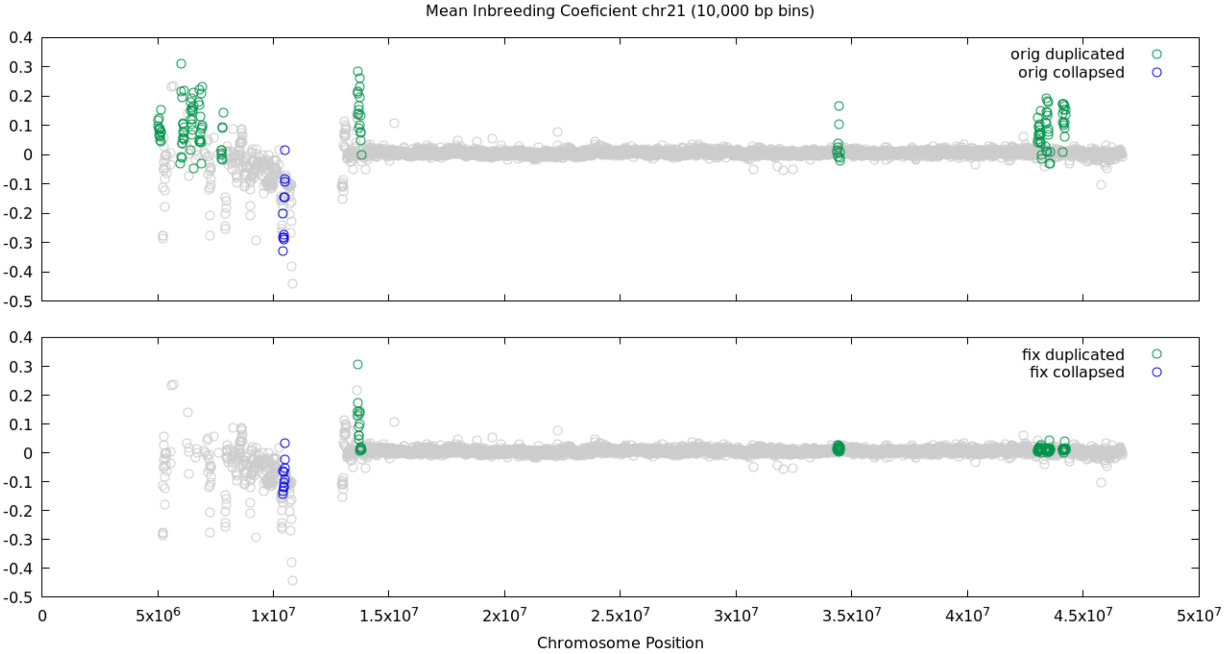


***Supplementary Figure S3.n: Binned inbreeding coefficient.*** *Mean inbreeding coefficients for 10,000 basepair bins of chromosome 21.*

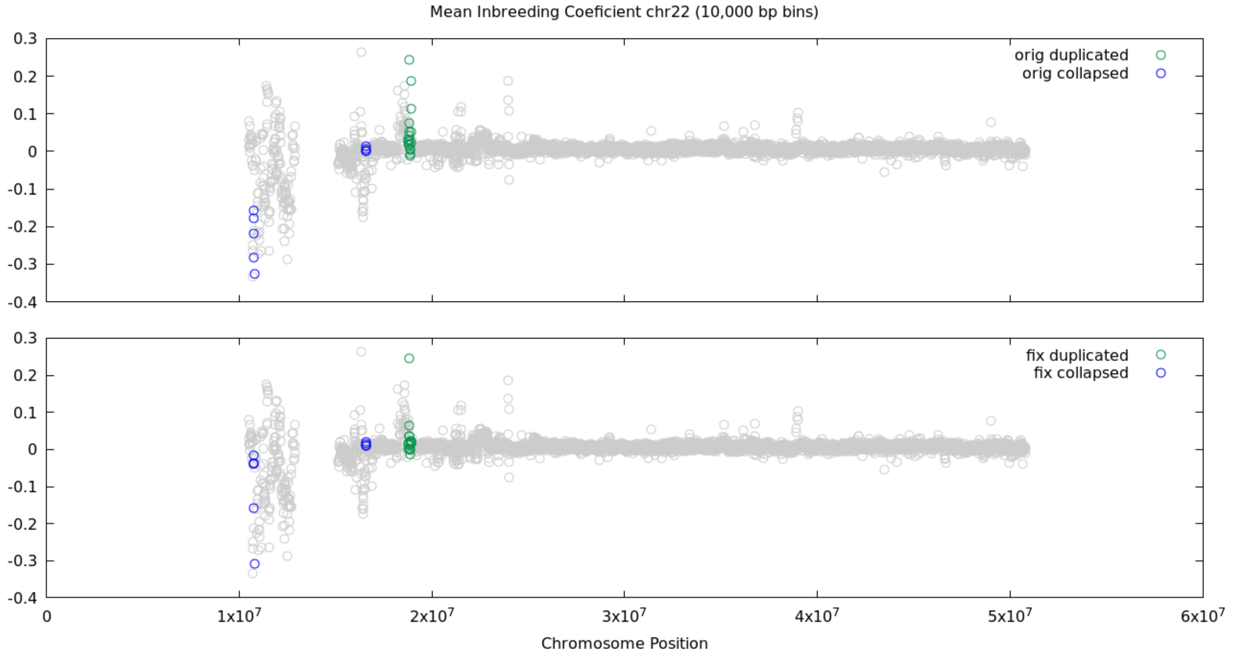


***Supplementary Figure S3.o: Binned inbreeding coefficient.*** *Mean inbreeding coefficients for 10,000 basepair bins of chromosome 22.*

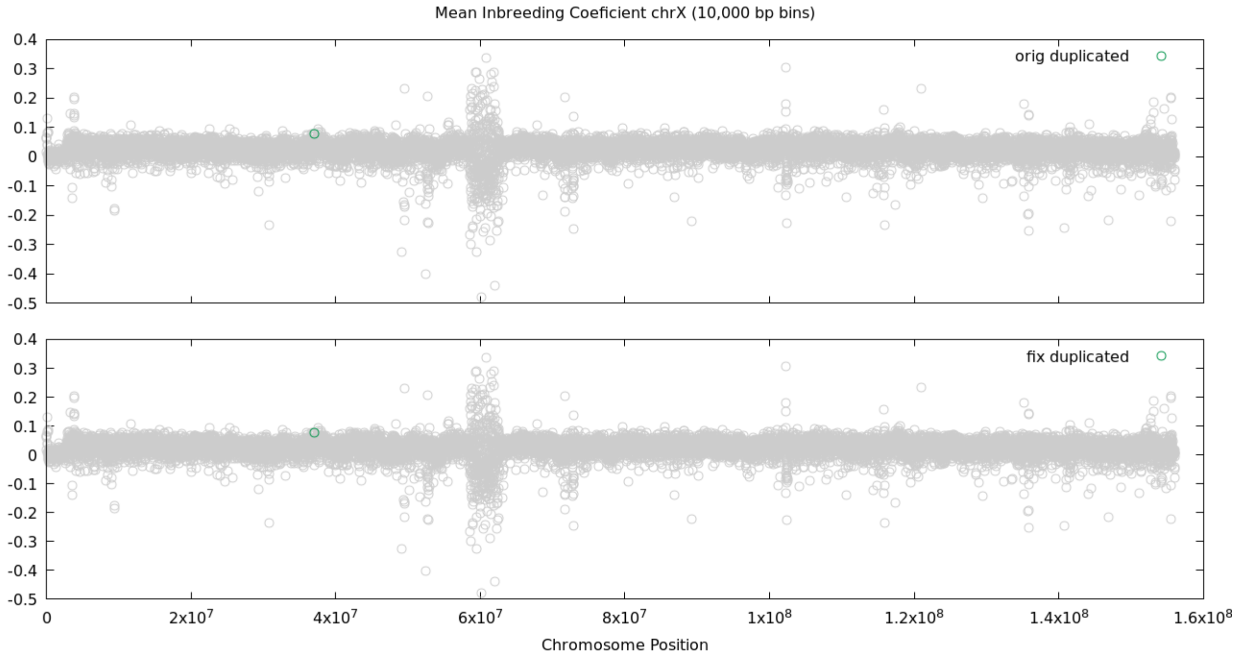


***Supplementary Figure S3.p: Binned inbreeding coefficient.*** *Mean inbreeding coefficients for 10,000 basepair bins of chromosome X.*

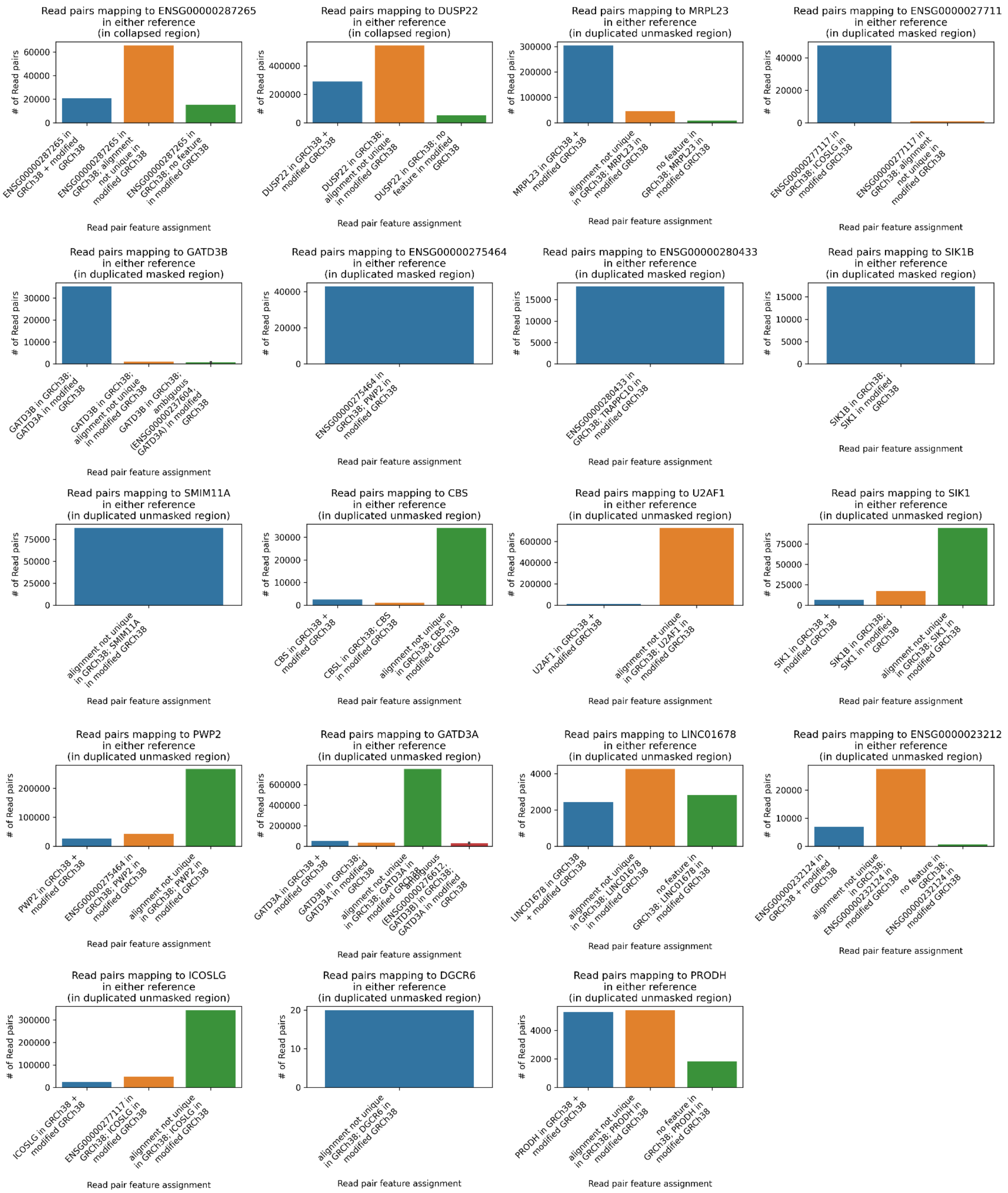


***Supplementary Figure S4: Comparison of RNA-seq read pair - gene assignments for reads mapping to the specified gene in at least one of the references.*** *Genes shown are those in collapsed, duplicated masked, or duplicated unmasked regions that have mean CPM >= 1 in at least one of the references and % change in CPM between the references of at least 10%. To maintain readability, features assignment combinations (x-axis categories) accounting for < 1% of the data in any panel are not displayed*

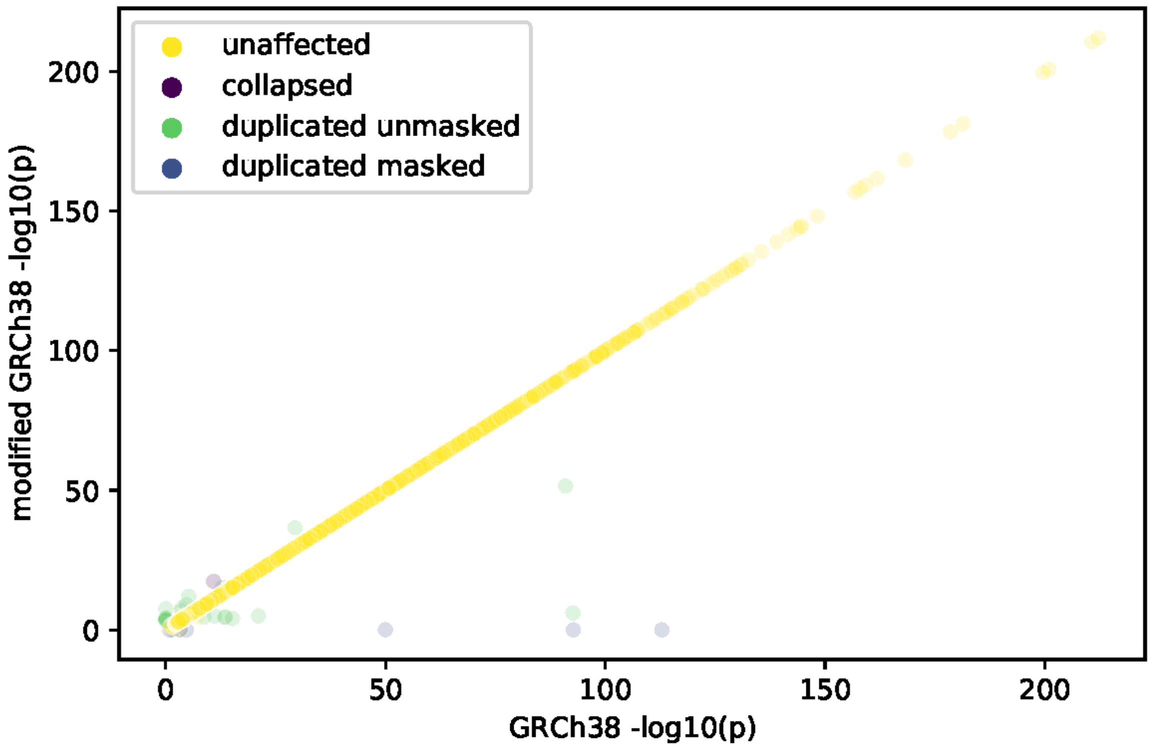


***Supplementary Figure S5:* Comparison of cis-eQTL p-values for genetic variants most significantly associated with each gene's expression in GRCh38 or the modified GRCh38**. *Genes are labeled according to whether or not they overlap collapsed, duplicated masked, or duplicated unmasked regions, or none of the above ('unaffected'). If a gene was not included in the eQTL scan for one of the references (because the expression was too low), the p-value was set to 1 for the purposes of this panel.*

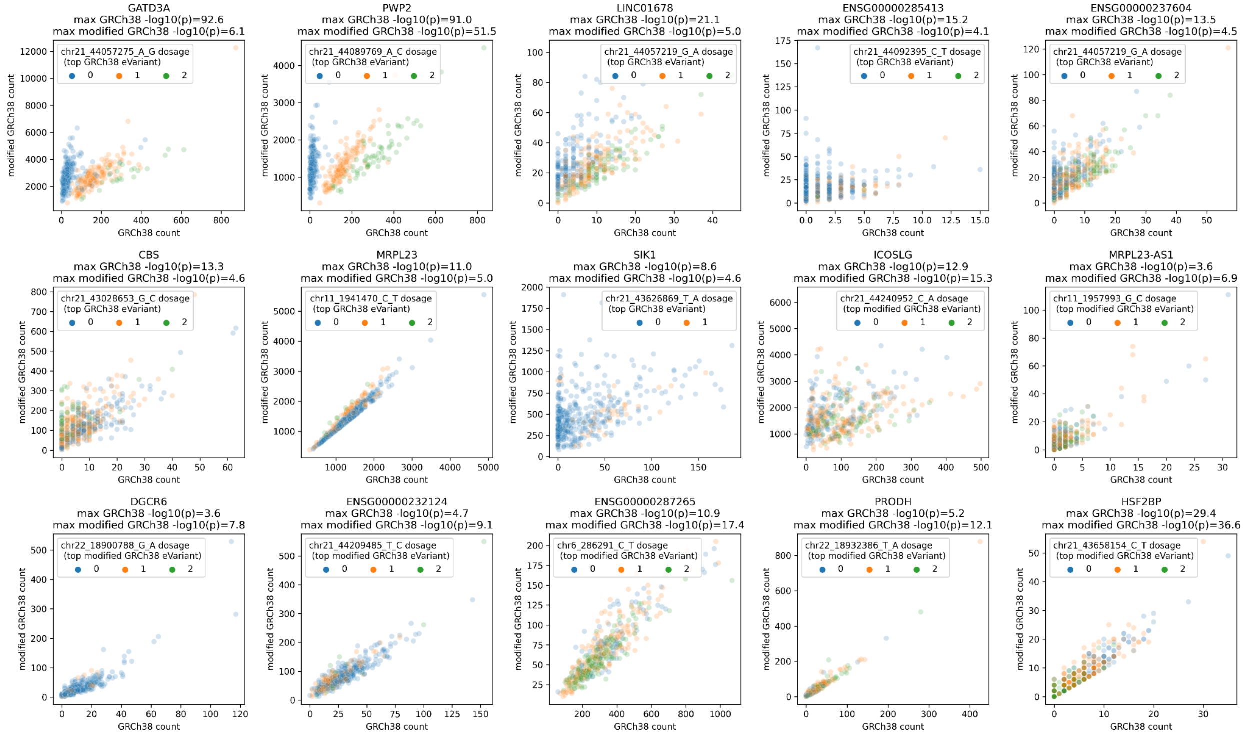


***Supplementary Figure S6:*** ***Per-sample gene read counts for the indicated gene when using GRCh38 or the modified GRCh38,*** *for genes meeting the following criteria: (1) tested in the eQTL scan for both references, (2) eGene in at least one of the references, (3) overlaps in collapsed/duplicated masked/duplicated unmasked regions, (4) -log10(p) value is at least one order of magnitude greater in one reference relative to the other. Samples are colored according to their genotype for the top gene expression associated genetic variant in the reference with the most extreme eQTL p-value for the gene*

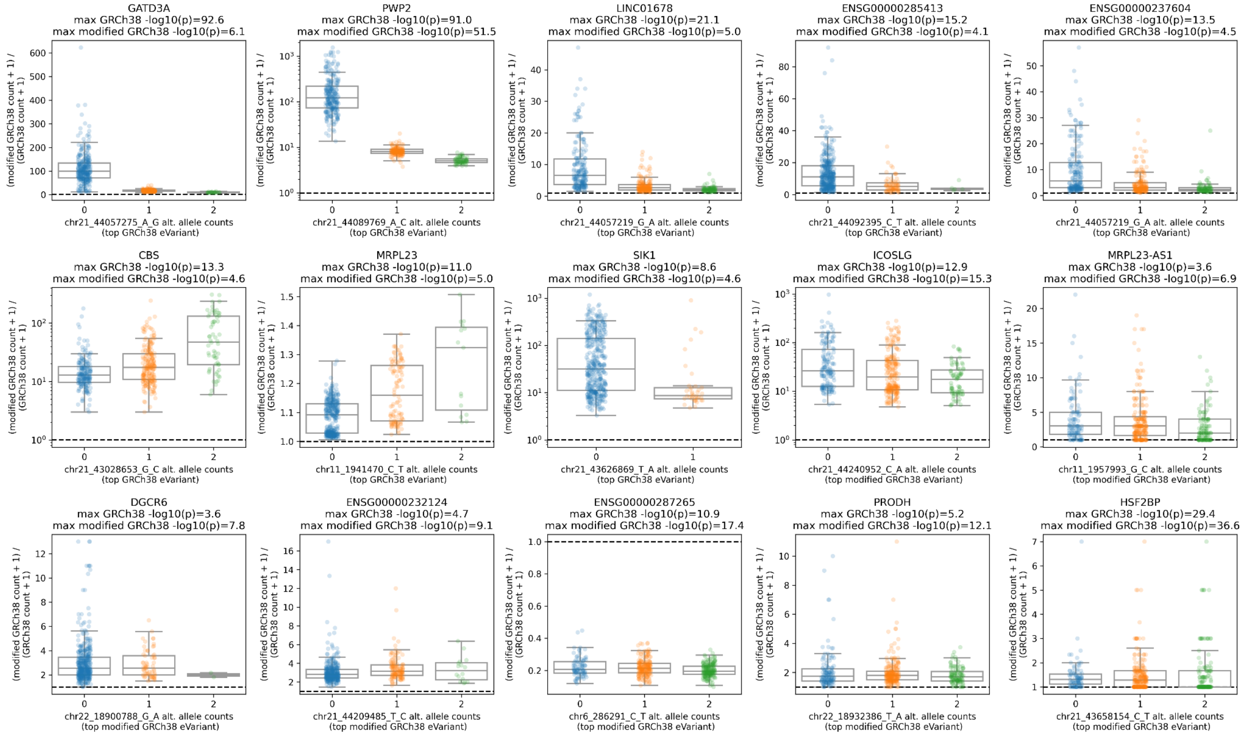


***Supplementary Figure S7:*** ***Per-sample ratio between gene read counts (+ pseudocount) for the indicated gene when using modified GRCh38 or GRCh38 reference to quantify gene expression***. *Samples are stratified and colored according to their genotype for the top gene expression associated genetic variant in the reference with the most extreme eQTL p-value for the gene. Genes shown meet the following criteria: (1) tested in the eQTL scan for both references, (2) eGene in at least one of the references, (3) overlaps in collapsed/duplicated masked/duplicated unmasked regions, (4) -log10(p) value is at least one order of magnitude greater in one reference relative to the other. Genes with stronger max eQTL p-values in GRCh38 than in modified GRCh38 display genotype-associated differences in % change in expression between the two references, consistent with genotype-related mapping biases. Such strong genotype-associated differences are generally not apparent for genes with stronger max eQTL p-values in GRCh38.*

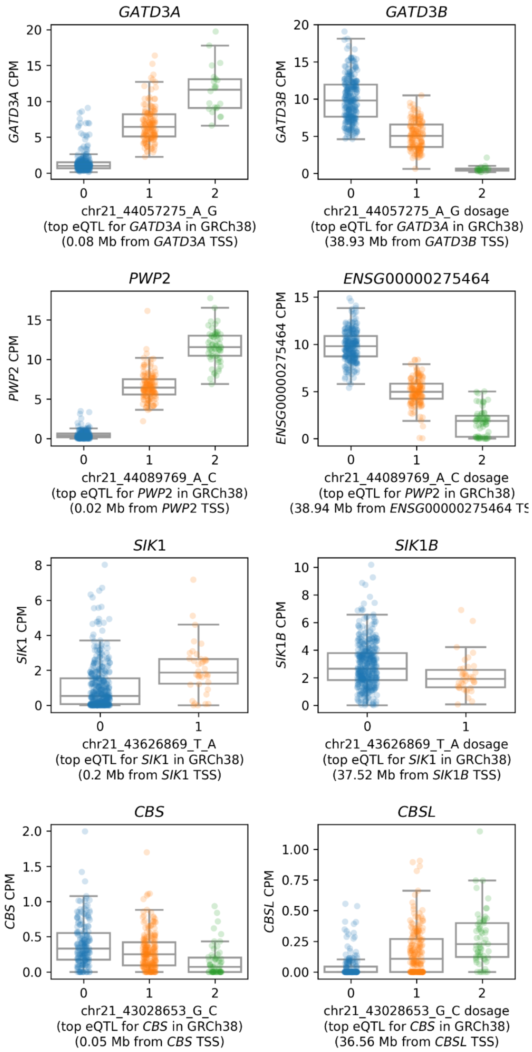


***Supplementary Figure S8:***

*For a subset of genes in falsely duplicated regions, the variant most strongly associated with the falsely duplicated genes in GRCh38 is also associated with the false copy of the same gene, with the opposite direction of effect. Each row represents one pair of duplicated genes; the genes in the left column are the true copies, while the genes in the right column are the false copies. X-axis values represent the genetic variant alternate allele dosage for the variant most strongly associated with the true copy; y-axis values represent gene expression (normalized to counts per million).*

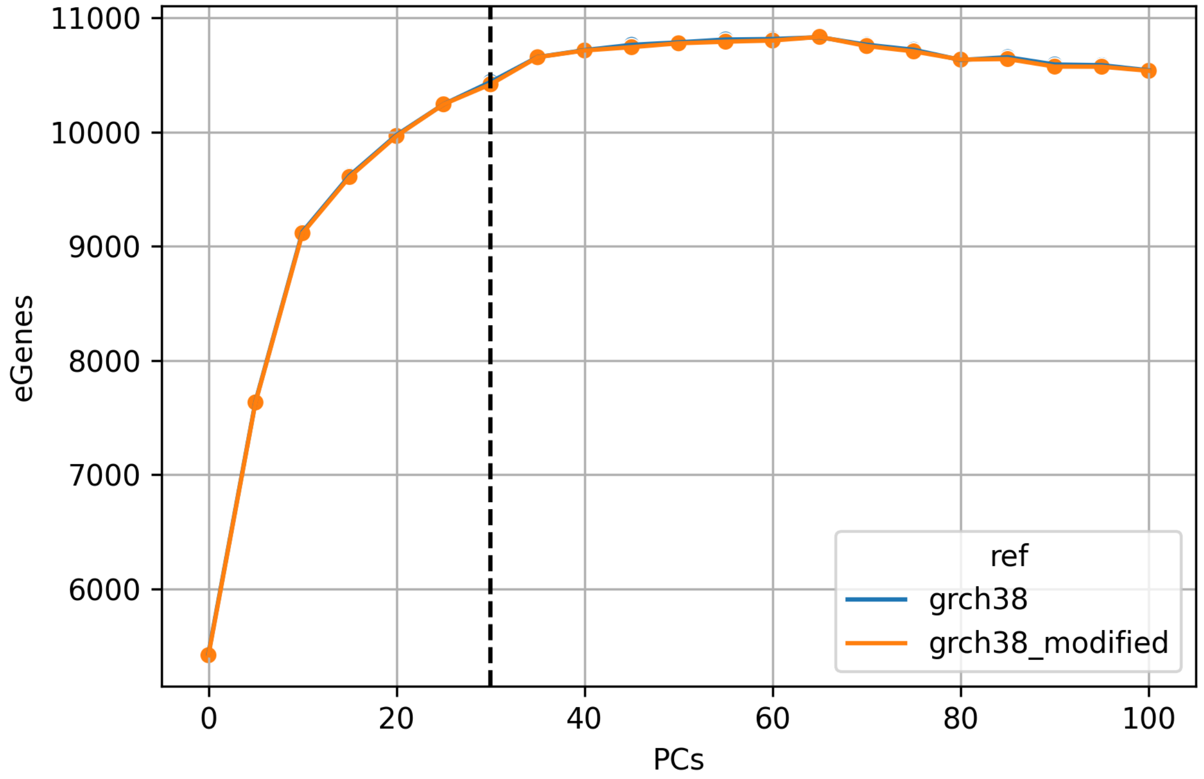


***Supplementary Figure S9:***

*Number of eGenes discovered (y-axis) vs number of gene expression PCs included in the eQTL scan (x-axis). The vertical line at x = 30 PCs represents the number of PCs used in the final eQTL scans.*
